# Separate lanes for math and reading in the white matter highways of the human brain

**DOI:** 10.1101/420216

**Authors:** Mareike Grotheer, Zonglei Zhen, Garikoitz Lerma-Usabiaga, Kalanit Grill-Spector

**Affiliations:** Psychology Department, Stanford University, Stanford, USA; Beijing Key Laboratory of Applied Experimental Psychology, Faculty of Psychology, Beijing Normal University, Beijing, China; Stanford Neurosciences Institute, Stanford University, Stanford, USA

**Keywords:** Reading, math, language, network, connectome, anatomy, functional MR, diffusion MR

## Abstract

Math and reading involve distributed brain networks and have both shared (e.g. encoding of visual stimuli) and dissociated (e.g. quantity processing) cognitive components. To date, it is unknown what are shared vs. dissociated gray and white matter substrates of the math and reading networks. Here we address this question using an innovative, multimodal approach applying functional MRI, diffusion MRI, and quantitative MRI to define these networks and evaluate the structural properties of their fascicles. Results reveal that i) there are distinct gray matter regions which are preferentially engaged in either math or reading and ii) the superior longitudinal (SLF) and arcuate (AF) fascicles are shared across math and reading networks. Strikingly, within these fascicles, reading- and math-related tracts are segregated into parallel sub-bundles and show structural differences related to myelination. These novel findings open a new avenue of research that examines the contribution of sub-bundles within fascicles to specific behaviors.

---

Math and reading are essential for functioning in modern society. We are not born with these skills, but rather acquire them through extensive learning, typically in childhood. While math and reading are distinct tasks, they utilize several overlapping cognitive processes, including encoding of visual stimuli, verbalization, as well as working memory^1^. It has been suggested that the degree to which brain activations related to math and reading overlap may depend on the arithmetic ask. For example, brain responses related to arithmetic fact retrieval (e.g. during addition of small numbers) overlap more with brain responses related to reading than responses related to procedural-based computations^2^. There is also a surprisingly high rate of co-morbidity between math and reading disabilities: up to 66% of children affected by dyscalculia, a math learning disability, also suffer from dyslexia, a reading learning disability (for review see^3^). Together with the shared cognitive components, this high rate of comorbidity suggests that math and reading may rely on shared neural substrates. While a large body of research has examined both cortical regions and white matter connections of the reading network^4,5^, the cortical regions and white matter connections of the math network as well as how they relate to the reading network are not well understood (see recent reviews^6,7^).

Research indicates that several white matter fascicles are key for reading. The fascicles of the reading network include: (i) The arcuate fasciculus (AF), which connects the frontal and temporal cortices. Diffusion MRI (dMRI) measurements show that fractional anisotropy (FA) in the left AF correlates with phonological awareness in both typical^8^ and impaired^9,10^ readers and that dyslexics have reduced FA in the left AF^9,11^. (ii) The inferior fronto-occipital fasciculus (IFOF), which connects the frontal and occipital cortices. Children with dyslexia show a reduced leftward asymmetry of the IFOF^12^ and FA of this tract is linked to orthographic processing skill^9,13^. (iii) The inferior longitudinal fasciculus (ILF), which connects the occipital lobe with the anterior tip of the temporal lobe. Lesions to the ILF can lead to pure alexia^14^, and atypical development of FA in the ILF is associated with poor reading proficiency^15^ as well as dyslexia^10^. (iv) The vertical occipital fasciculus (VOF), which connects the occipital and parietal cortices^16^ and is thought to relay top-down signals from the intraparietal sulcus (IPS) to ventral occipito-temporal cortex during reading^17^. Interestingly, these four fascicles intersect with the visual word form area (VWFA)^18–20^, a region in the occipito-temporal sulcus (OTS) that responds preferentially to words over other stimuli. The VWFA is thought to process visually presented words^21,22^ and is causally involved in word recognition, as lesioning it produces acquired dyslexia^23^.

Although it has been suggested that math and reading utilize several overlapping cognitive processes^1^, presently it is unclear if fascicles associated with reading are also associated with mathematical processing. This fundamental gap in knowledge is due to four main reasons: First, substantially more neuroscience research has been done on the neural bases of reading^4,5^ than the neural bases of math^6^. Second, most prior studies have evaluated either the neural bases of math^7,24–27^ or the neural bases reading^18,21,28–33^, but not both systems within the same individuals (for an exception see^2^). Third, no study directly examined the white matter connections associated with functional regions involved in mathematical processing within the same individuals. Previous work on the neural substrates of mathematical processing has focused on linking gray matter structure, evaluated with voxel-based morphometry, and white matter (^34^, for review see^6,7^). The only study that evaluated the white matter connections of functional regions related to mathematical processing evaluated gray and white matter in distinct groups of participants^35^. While this study provided first insight on the white matter connections of the math network, more work is needed to understand these connections within the same individuals. In contrast, a few previous studies have examined which white matter fascicles connect to a key cortical region of the reading network, namely the VWFA, evaluating functional regions and white matter connections within the same subjects^18–20^. Thus, the goal of this study is twofold: (1) identify and quantify the white matter connections of cortical regions involved in mathematical processing within individual subjects and (2) determine which aspects of this white matter are unique to the math network, and which are shared with the reading network.

To fill these gaps in knowledge we applied an innovative, multimodal approach, in which we collected functional MRI (fMRI), diffusion MRI, and quantitative MRI (qMRI) data in 20 participants. The goal of the fMRI experiment was to identify in each participant’s brain the cortical regions that are involved in reading, mathematical processing, or both. As in our prior study^36^, we presented participants with number-letter morphs, such that the three tasks, reading, adding or remembering colors, could be performed on identical stimuli (**Fig. 1a**). The goal of the dMRI measurements was to determine the white matter tracts of the math and reading networks. Using dMRI data and modern tractography methods that account for crossing fibers^37–39^, we generated a white matter connectome in each participant and then, using Automated Fiber Quantification (AFQ)^40^, identified 13 well-established fascicles in each participant, most of them bilaterally. Critically, to identify functionally-defined white matter tracts (fWMT) of the math and reading networks, first we identified the gray-white matter interface (GWMI) of functional regions of interest (fROIs) involved in reading, adding, or both. We then determined which white matter fiber tracts are seeded or terminate in the GWMI of each fROI (**Fig. 1c**), as well as, which fascicles these fWMTs travel through. This approach allowed us to determine (1) what are the white matter tracts of the math and reading networks, (2) which fascicles are network unique and which contribute to both math and reading, and (3) whether white matter tracts associated with math and reading are physically intertwined or segregated within a fascicle. Finally, the goal of the qMRI measurements was to test if there are differences in structural white matter characteristics across the reading and math networks. qMRI^41–43^ allowed us to measure the proton relaxation time (T_1_) of the identified fWMT of each network. As T_1_ in the white matter is correlated with myelination^44^, changes with development and varies across fascicles^45^, it enabled *in vivo* assessment of microstructural properties of the identified tracts.

Our multimodal approach thus provides a unique opportunity to determine what are the shared and segregated white matter tracts of the math and reading networks, as well as measure their structural properties. Thereby our approach broadens the current understanding of the brain substrates of fundamental skills learned by children world-wide, which, in turn, may provide a key stepping stone for understanding how childhood education shapes the brain.

## Results

### Neighboring gray matter regions process math and reading

We first used fMRI to define gray matter regions that are involved in math and reading in each participant. In the fMRI experiment, participants performed a reading task, a math task, and a color memory task on identical visual stimuli (number-letter morphs, **Fig. 1a**). In each trial, subjects viewed a cue indicating the task (“Read”/”Add”/”Color”), then viewed four number-letter morph stimuli that were presented sequentially, and at end of the trial gave a task-relevant 2-alternative-forced choice answer. In the math (adding) task, subjects summed up the presented stimuli and indicated which of two numbers shows the correct sum. In the reading task, they were instructed to read the word and indicate which of the two words matches the one they had read, and in the color task they were instructed to attend to the color of the stimuli and indicate which of 2 colors had been used for one of the presented characters. Crucially, these tasks were performed on identical visual stimuli and were matched in their working memory load and the amount of verbalization they elicit.

Participants successfully performed all tasks in the experiment (average accuracy (±SE): 88.16(2.43)%). Both accuracy and response times (RTs) differed across the reading, adding, and color tasks (main effect of task: accuracy: F(2,38) = 10.30, p=0.0003, ηp^2^=0.35; RTs: F(2,38)=72.20, p<0.0001, ηp^2^=0.79). While accuracy was significantly higher in the reading task, relative to the other two tasks (all ps<0.002 after Bonferroni correction, n.s. between adding and color), response times were shortest in the adding, intermediate in the reading task, and slowest in the color task (all ps<0.003 after Bonferroni correction). It is unlikely that performance differences across tasks drove responses across cortex for two reasons: i) within a task, response accuracy and neural task preference, i.e. the extent of preferential activations for a given task, did not show any clear relationship across participants (parameter maps presented in **Supplementary Fig. 1-4** are sorted according to task performance, for group maps see **Supplementary Fig. 5**), and ii) accuracy and response times were not consistently different across tasks, yet we could identify task-selective functional regions of interest (fROIs) for all tasks (**Fig. 1b, Supplementary Fig. 6-7**).

Regions involved in math were defined by higher responses during the math task than the reading and color tasks, while regions involved in reading were defined by higher responses during the reading task than the math and color tasks (as in our prior study^36^). We also considered regions involved in both math and reading, defined by the conjunction of both higher responses during math than color as well as during reading than color task (conjunction analysis, math > color ∩ reading > color). Crucially, all regions were defined in individual subjects’ native anatomical space and without spatial smoothing, as both group averaging and spatial smoothing may introduce artificial overlap between regions^46^.

We found consistently stronger responses during the reading task compared to adding and color tasks in four anatomical expanses (example subject in **Fig. 1b-green**; all subjects in **Supplementary Fig. 6-7**): (i) A region in the occipito-temporal sulcus (OTS) (left hemisphere: N=18, size±SE: 492±97 mm^3^; right hemisphere: N=12, size ±SE: 107±33 mm^3^). Activations in the OTS were frequently divided into two distinct subregions and likely correspond to the visual word form areas (VWFA1 and VWFA2^21,22^). Here, we took their union as a single fROI, as we were interested in determining the large-scale white matter networks associated with reading and math. (ii) A region in the superior temporal sulcus (STS), which extended into the middle temporal sulcus (left hemisphere: N=20, size±SE: 774±184 mm^3^; right hemisphere: N=17, size±SE: 343±110 mm^3^). (iii) A region in the supramarginal gyrus (SMG, we will refer to this region in the reading network as SMGr) (left hemisphere: N=20, size±SE: 494±148 mm^3^; right hemisphere: N=19, size±SE: 119±34 mm^3^). (iv) A region in the inferior frontal gyrus (IFG), which likely corresponds to “Broca’s Area” (left hemisphere: N=20, size±SE: 1458±255 mm^3^; right hemisphere: N=20, size±SE: 668±157 mm^3^). Activations in the IFG spanned 2-3 clusters and we took their union. Activations during reading were substantially smaller and less frequent in the right hemisphere than the left hemisphere, thus we focus on the left hemisphere in the main document and present right hemisphere data in the Supplementary Material (**Supplementary Fig. 10, 12, 17, 21**).

We also identified four bilateral regions that responded more strongly during the adding than reading or color tasks (example subject in **Fig. 1b-blue**; all subjects in **Supplementary Fig. 6-7**): (i) A region in the inferior temporal gyrus (ITG; left hemisphere: N=20, size±SE: 680±126 mm^3^; right hemisphere: N=19, size±SE: 570±92 mm^3^), which is consistent with prior studies^36,47^. (ii) A region in the intra-parietal sulcus (IPS; left hemisphere: N=19, size±SE: 1329±224 mm^3^; right hemisphere: N=18, size±SE: 1283±197 mm^3^), which is in line with previous research showing IPS involvement in numerosity processing^24,26^. (iii) A region in the SMG (we will refer to this region in the math network as SMGm) (left hemisphere: N=20, size±SE: 893±178 mm^3^; right hemisphere: N=20, size±SE: 1170±187mm^3^). (iv) A region in the inferior part of the precentral sulcus (PCS; left hemisphere: N=18, size±SE: 763±141 mm^3^; right hemisphere: N=19, size±SE: 580±100 mm^3^). This region is anatomically proximal to the inferior frontal junction, which has been implicated in visual object-based attention^48^.

**Figure 1.**
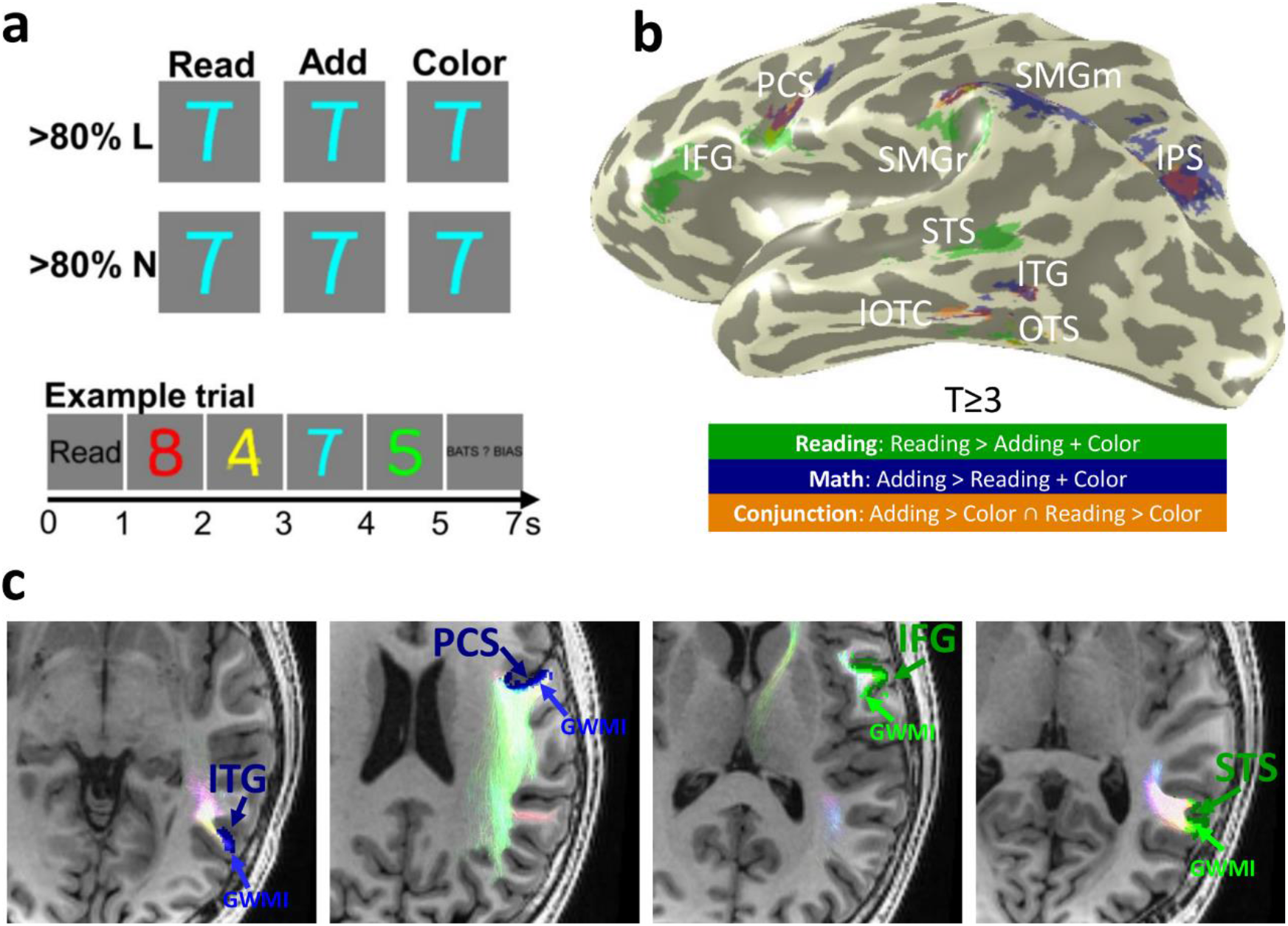
Identification of gray matter regions of the math and reading networks and their fWMT. **(a)** FMRI experiment used to define math- and reading-related regions. Subjects viewed morphs between numbers and letters, containing either >80% letter (<20% number) or >80% number (<20% letter) information. At the beginning of each trial, a cue (“Read”/”Add”/”Color”) indicated which task should be performed, then 4 stimuli of the same morph type appeared for 1 sec each, followed by an answer screen presented for 2 secs. Subjects indicated their answer with a button press. Identical stimuli were presented across tasks. Trial structure is shown at the bottom. **(b)** Gray matter regions of the math and reading networks. *Green:* Reading-related regions were defined based on higher responses in the reading task than other tasks; *Blue:* Math-related regions were defined based on higher responses in the adding task than other tasks; *Orange:* Regions that responded more strongly during reading vs color and adding vs color tasks. All fROIs were defined using a T ≥ 3 (voxel level) threshold in each participant’s brain. **(c)** Example fROIs and their respective fWMTs in a representative participant’s axial slices. *Blue:* Math fROIs. *Green:* Reading fROIs; lighter shades of blue and green under each fROI: respective GWMI of that fROI. The fiber tracts that terminate at the GWMI of each fROI in this slice are shown in pastel colors; the colors of the tracts indicate the main diffusion direction (pink: right/left; light green: anterior/posterior; light blue: superior/inferior). *Abbreviations:* IFG=inferior frontal gyrus, PCS=precentral sulcus, SMGr=reading fROI in supramarginal gyrus, SMGm=math fROI in supramarginal gyrus, STS=superior temporal sulcus, ITG=inferior temporal gyrus, OTS=occipito-temporal sulcus, IPS=intraparietal sulcus, lOTC=lateral occipito-temporal cortex, fWMT=functionally-defined white matter tracts, GWMI=gray/white matter interface.

Interestingly, gray matter regions involved in reading and math were often neighboring. In the prefrontal cortex, the reading-related IFG is proximal, but inferior to the math-related PCS. Likewise, in the SMG the reading-related fROI (SMGr) was proximal, but inferior to the math-related fROI (SMGm). In the temporal cortex, math-related ITG is located between two reading-related fROIs, centered on the STS and OTS. In the IPS we found only a math-related fROI.

Four regions in the brain showed higher responses during both the adding and reading tasks compared to the color task (conjunction analysis, math > color ∩ reading > color; **Fig. 1b-orange, Supplementary Fig. 6-7**): (i) A region in the intra-parietal sulcus (IPS; left hemisphere: N=18, size±SE: 363±90 mm^3^; right hemisphere: N=16, size±SE: 251±63 mm^3^). (ii) A region in the SMG (SMG, left hemisphere: N=20, size±SE: 434±89 mm^3^; right hemisphere: N=17, size±SE: 249±54 mm^3^). (iii) A region in the inferior part of the precentral sulcus (PCS; left hemisphere: N=19, size±SE: 350±87 mm^3^; right hemisphere: N=18, size±SE: 111±26 mm^3^). (iv) A region in the lateral occipitotemporal cortex (lOTC) that extended from the ITG to the OTS (left hemisphere: N=19, size±SE: 720±141 mm^3^; right hemisphere: N=18, size±SE: 241±53 mm^3^). Except for the lOTC region, these fROIs were small and tended to overlap with the math fROIs (**Fig 1b, Supplementary Fig. 6-7**). Indeed, responses in the IPS, PCS, and SMG conjunction fROIs were significantly stronger during the adding task than the reading task (**Supplementary Fig. 8**; ANOVA with hemisphere, task and stimulus as factors; main effect of task: IPS: F(1,14) = 17.30, p=0.001, ηp^2^=0.55; PCS: F(1,16) = 12.97, p=0.002, ηp^2^=0.45; SMG: F(1,16) = 19.37, p=0.0004, ηp^2^=0.55). The only conjunction fROI in which responses during adding and reading did not differ significantly was the lOTC (main effect of task: F(1,17)=3.83, p=0.07, ηp^2^=0.18). This region overlaps with regions in both the math network (in the ITG) and the reading network (in the OTS). Thus, in subsequent analyses we considered the IPS, PCS, and SMG regions as part of the math network, but the lOTC as a conjunction region involved in both tasks.

### The SLF and the AF contribute to math and reading networks

After establishing which cortical regions are activated during math and/or reading tasks, we determined which fascicles are associated with each of these fROIs (**Fig. 2**). For this, we identified 13 well-established fascicles of the brain, most of them bilaterally, in each participant using AFQ (for visualization of all fascicles see **Supplementary Fig. 9**). Next, we intersected each participant’s classified white matter connectome with the GMWI underneath each of the fROIs to determine the functionally-defined white matter tracts (fWMT) associated with reading and/or math (**Fig. 1c**). To summarize the fascicles connecting to each reading and math related region across subjects, we quantified the percentage of the fWMT associated with each of the 13 fascicles, separately for each fROI. We will refer to the relative contribution of each fascicle as “connectivity weight”. Results reveal two main findings.

First, across participants, six fascicles contain almost all the fWMT of math and reading fROIs (sum of their connectivity weights is above 90%). These fascicles are: inferior fronto-occipital fasciculus (IFOF), inferior longitudinal fasciculus (ILF), superior longitudinal fasciculus (SLF), arcuate fasciculus (AF), posterior AF (pAF), and vertical occipital fasciculus (VOF). In **Fig. 2**, the left panels show the fWMT of math, reading, and the lOTC conjunction fROIs in a representative subject, the middle panels show the connectivity weight of these fascicles across subjects, and the right panels provide a schematic illustration of the same data, depicting the general anatomical layout. Notably, three out of these six fascicles, the SLF, AF and pAF, form the backbone of the math and reading networks. For at least one fROI in each network, these fascicles contain >10% of all fWMT. A similar pattern of results is observed in the right hemisphere (**Supplementary Fig. 10**), and when we controlled for fROI size, by repeating the analyses using constant-size spherical ROIs (radius=7mm) centered on the fROIs (**Supplementary Fig. 11**).

**Figure 2.**
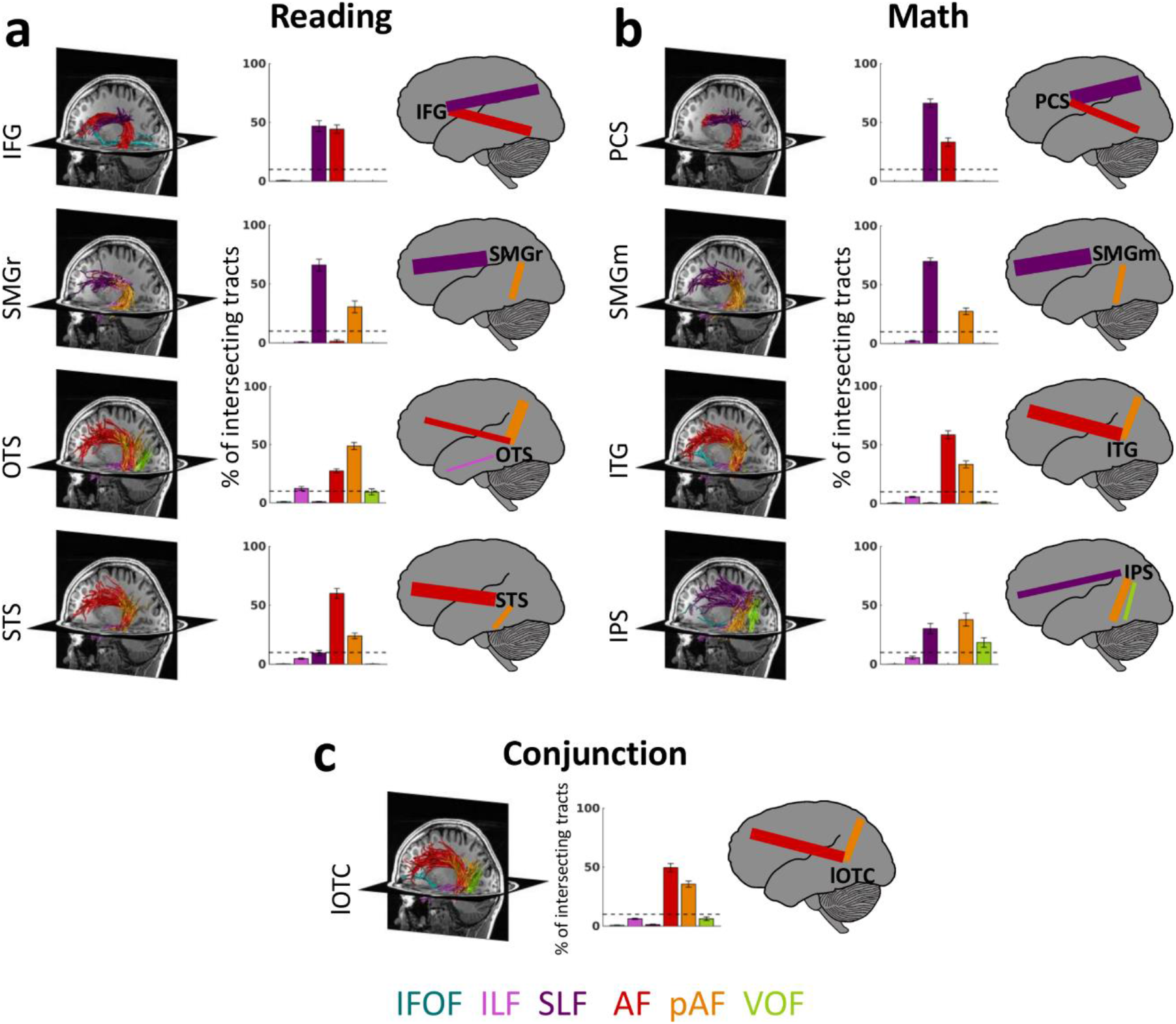
Functionally-defined white matter tracts (fWMT) of reading and math related regions. **(a)** Six fascicles (AF, SLF, pAF, VOF, ILF, and IFOF) contain >90% of all fWMT of the fROIs identified in the reading task. **(b)** The same six fascicles also contain >90% of all fWMT of the fROIs identified in the math task. **(c)** The conjunction fROI in the lOTC shows substantial connectivity with the AF and pAF. In **(a, b, c)**: *Left:* fWMT for each fROI in a representative subject’s left hemisphere. The same subject is displayed in all panels; Fascicles are color coded in accordance with the legend at the bottom. *Middle.* Bar graphs showing what percentage of the fWMT is associated with each of the six fascicles. The graph shows the mean across subjects ± SEM. *Dashed horizontal line:* Line is placed at 10%, which was the cut-off used for the schematics in the right columns. *Right:* Schematic illustration of the fascicles associated with each fROI. The thickness of the lines is derived from the bar graph, showing the relative weight of each fascicle. *Abbreviations:* IFG=inferior frontal gyrus, PCS=precentral sulcus, SMGr=reading fROI in supramarginal gyrus, SMGm=math fROI in supramarginal gyrus, STS=superior temporal sulcus, ITG=inferior temporal gyrus, OTS=occipito-temporal sulcus, IPS=intraparietal sulcus, lOTC=lateral occipito-temporal cortex, IFOF=inferior fronto-occipital fasciculus, ILF=inferior longitudinal fasciculus, SLF=superior longitudinal fasciculus, AF=arcuate fasciculus, pAF=posterior arcuate fasciculus, VOF=vertical occipital fasciculus.

Second, we found that anatomically neighboring math and reading fROIs in the prefrontal cortex and SMG connect to the same fascicles with a comparable weight. That is, they show a relatively similar connectivity fingerprint. This is evident for the reading fROI in the IFG and the math fROI in the PCS that illustrate (i) substantial connectivity to the SLF (connection weight > 46%), which connects the frontal and parietal lobes, (ii) substantial connectivity to the AF (connection weight > 33%), which connects the fontal and temporal lobes, and (iii) no substantial connections to other fascicles (**Fig. 2**, first row). Similarly, both the reading and math fROIs in the SMG (SMGr and SMGm, respectively) are: (i) strongly connected to the SLF (weight >66%), (ii) connected to the pAF (> 27%), which connects the parietal and temporal lobes, and (iii) show no substantial connections to other fascicles (**Fig. 2**, second row). In comparison, reading and math fROIs in the temporal cortex and IPS showed more differentiated connections across networks. For example, while the reading fROI in the OTS showed above 10% connection weight with the AF, pAF, and ILF, a nearby math fROI in the ITG showed above 10% connection weight for the former two, but not the latter (**Fig. 2**, third row). Finally, the lOTC conjunction fROI was mainly connected to the AF (49.57%) and pAF (35.60%) (**Fig. 2c**).

Next, we determined within-network connections of the math and reading networks. To identify all pairwise connections, we intersected the fWMT of each fROI with the GWMWI of each of the other non-neighboring fROIs within each network (**Fig. 3a,e** shows a representative subject). We quantified the pairwise connection relative to the total fWMT of each of the fROIs constituting the pair using the dice coefficient (DC^49^), and evaluated if it was significantly greater than chance level (**Fig. 3b,f**) . The DC indicates the proportion of fWMT shared between two fROIs relative to the total fWMT of these fROIs.

In the reading network, we find significantly above chance DC (**Fig. 3a,b**; significance was determined with a Bonferroni-adjusted threshold of p<0.008) between: (i) the OTS and the STS (paired t-test: p=0.0004, t(17)=4.4), (ii) the OTS and the IFG (paired t-test: p<0.0001, t(17)=5.55), (iii) the STS and the IFG (paired t-test: p=0.0003, t(19)=4.42), and (iv) SMGr and the IFG (paired t-test: p<0.0001, t(19) = 5.07). The significant frontal-temporal connections of the reading network (STS-IFG; OTS-IFG) are supported by the AF, the frontal-parietal connections (IFG-SMGr) are supported by the SLF and the ventral-temporal connections (OTS-STS) are supported by the pAF (**Fig. 3c**).

In the math network, we find significantly above chance DC (**Fig. 3e,f** significance was determined with a Bonferroni-adjusted threshold of p<0.008) between (i) the ITG and SMGm (paired t-test: p=0.005, t(19)=3.18, **Fig. 3f**), which is supported by the pAF (**Fig. 3g**), between (ii) the ITG and the PCS (paired t-test: p=0.0002, t(17)=4.77, **Fig. 3f**), via the AF (**Fig. 3g**), and (iii) between SMGm and the PCS (paired t-test: p=0.0003, t(11) = 5.26, **Fig. 3f**), through the SLF (**Fig. 3g**).

We summarize the pairwise connections and their predominant contributing fascicles in a schematic of within-network connections (**Fig. 3d**-reading, **Fig 3h**-math). Overall, these analyses suggest that both the math and the reading network illustrate significant within-network connectivity, and the AF and the SLF emerge as key fascicles in both networks.

We also evaluated between-network connectivity in two ways (i) by examining the pairwise connections of the conjunction lOTC fROI to each of the math and reading fROIs (**Fig. 3i-l**) and (ii) by evaluating the connections between pairs of fROIs across networks, where in each pair, one fROI was part of the reading network and the other fROI was part of the math network (**Fig. 3m-o**). Similar to the analyses of within-network connections described above, we only examined the long-range connections via fascicles, but not the local connections to neighboring fROIs.

For the conjunction fROI in the lOTC, when testing pairwise connections to fROIs of the reading network, we found significantly above chance DC (**Fig. 3i-l**, Bonferroni-adjusted threshold of p<0.01) only for connections to the IFG (paired t-test: p<0.0001, t(18)=5.12, **Fig 3j**), which are supported by the AF (**Fig 3k**). In contrast, we found significantly above chance connections of the conjunction fROI in the lOTC to several fROIs in the math network (i) the PCS (paired t-test: p-0.0007, t(18)=4.08), via the AF (ii) the SMGm (paired t-test: p=0.002, t(18)=3.70) via the pAF and (iii) the IPS (paired t-test: p=0.0002, t(16)=4.88) also via the pAF (**Fig. 3k**). Connections to both networks are summarized in a schematic (**Fig 3l**).

Analyzing pairwise connections across fROIs of the math and reading networks revealed significantly above chance DC (**Fig. 3n**, Bonferroni adjusted threshold of p<0.004) between: (i) the OTS and the IPS (paired t-test: p<0.0001, t(17)=5.10), (ii) the OTS and SMGm (paired t-test: p=0.0007, t(17)=4.14), (iii) the OTS and the PCS (paired t-test: p=0.003, t(15)=3.59), (iv) the STS and the PCS (paired t-test: p=0.001, t(17)=3.96), (v) SMGr and the PCS (paired t-test: p=0.0002, t(17)=4.72), (vi) the IFG and the ITG (paired t-test: p<0.0001, t(19) = 5.40), and (vii) the IFG and SMGm (paired t-test: p<0.0001, t(19)=5.06). Similar to the within-network connections described above, the pAF supported temporal-parietal between-network connections (OTS-IPS and OTS-SMGm), the SLF supported frontal-parietal between-network connections (SMGr-PCS and SMGm-IFG), and the AF supported frontal-temporal between-network connections (OTS-PCS, ITG-IFG) (data not shown). These connections are summarized in a schematic (**Fig 3o**).

Analysis of the right hemisphere as well as a control analysis using constant-size spherical ROIs (radius of 7mm), centered on the fROIs, shows a similar pattern of results (**Supplementary Fig. 12-13**).

**Figure 3.**
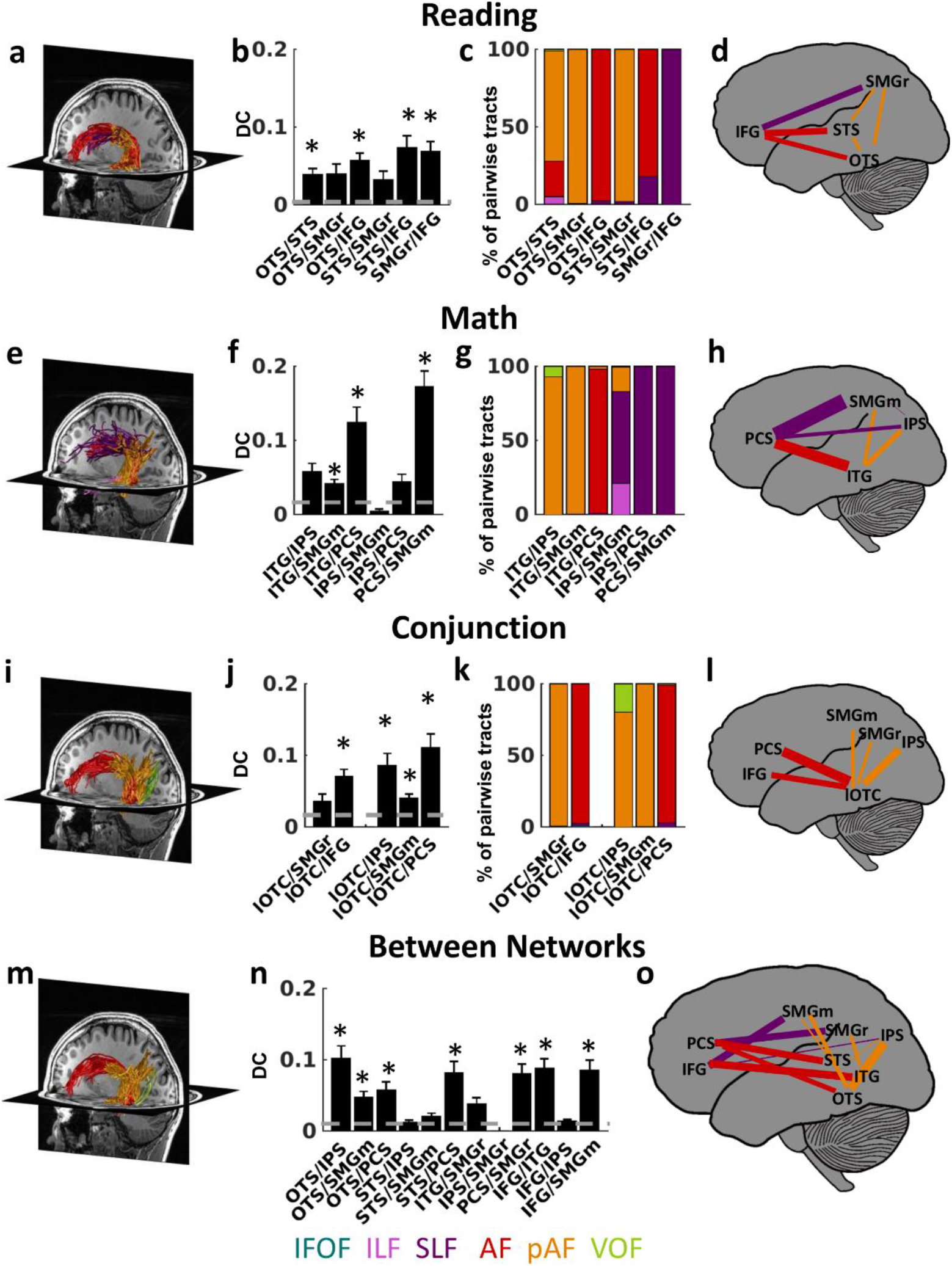
Pairwise fWMT within and between the reading and math networks. **(a-d)** Within-network connections of the reading network. **(e-h)** Within-network connections of the math network. **(i-j)** Connections of the conjunction fROI in the lOTC. **(m-o)** Between-network connections. *Left (a,e,i,m):* Pairwise white matter connections in a representative subject’s left hemisphere. *Second from left (b,f,j,n)*: Dice coefficient (DC) of pairwise connections, mean across subjects ± SEM. The DC quantifies the overlap in the fWMT of both fROIs: a DC of 1 indicates that all tracts that intersect with the first fROI also intersect with the second fROI, while a DC of 0 indicates no shared tracts. X-labels indicate the fROI pairing. *Dashed line:* Chance level DC estimated from the average connections to out of network fROIs in ventral temporal cortex that were activated maximally during the color task. *: DC is significantly higher than chance (significance level was Bonferroni adjusted). *Second from right in row 1-3 (c,g,k)*: The relative contribution of six fascicles to the pairwise connections (legend at bottom). X-labels indicate the fROI pairing. *Right (d,h,l,o):* Schematic illustration of the pairwise connections. *Line thickness* is scaled proportionally to the DC; *Color* indicates the fascicle with the highest relative contribution to pairwise connections. *Abbreviations:* IFG=inferior frontal gyrus, PCS=precentral sulcus, SMGr=reading fROI in supramarginal gyrus, SMGm=math fROI in supramarginal gyrus, STS=superior temporal sulcus, ITG=inferior temporal gyrus, OTS=occipito-temporal sulcus, IPS=intraparietal sulcus, lOTC=lateral occipito-temporal cortex, IFOF=inferior fronto-occipital fasciculus, ILF=inferior longitudinal fasciculus, SLF=superior longitudinal fasciculus, AF=arcuate fasciculus, pAF=posterior arcuate fasciculus, VOF=vertical occipital fasciculus.

### Reading and math tracts are segregated within SLF and AF

The analyses in the prior section highlight the AF and the SLF as the main fascicles of the reading network and the AF, SLF, and pAF as the main fascicles of the math network. The pAF and AF also contribute to between-network connection. Nonetheless, within-network pairwise connectivity (as quantified with the DC) was significantly higher than between-network connectivity (paired t-test comparing average within-network and between-network DCs: p=0.01, t(15)=2.83). Thus, in subsequent analyses we will focus on the within-network connections, as well as, on connection of the conjunction fROI in the lOTC to both the math and reading networks. We evaluated both the SLF and the AF, as these emerged as important fascicles of both the math and the reading network.

Since the SLF and AF are large and contain many tracts it is unclear whether these entire fascicles are part of both networks or, alternatively, if sub-bundles within these fascicles relay tracts that support within-network connectivity of the reading and math networks, respectively. We tested these hypotheses by visualizing and quantifying within the SLF tracts connecting the IFG and SMGr in the reading network as well as the PCS and SMGm in the math network, and by examining if they are spatially intertwined or segregated in each subject. Similarly, in the AF, we visualized and compared tracts connecting the PCS and the ITG in the math network, with those connecting the IFG and STS in the reading network. Across subjects, and in both the SLF and the AF, our data showed that tracts were segregated by network: fWMT supporting within-network connections of the math network (blue in **Fig. 4a,d**; for all subjects see **Supplementary Fig. 14-16**) were consistently superior to fWMT supporting within-network connections of the reading network (green in **Fig. 4a,d**; for all subjects see **Supplementary Fig. 14-16**).

Interestingly, in the AF, the connections of the lOTC conjunction fROI to the IFG (reading network) and the PCS (math network) were also segregated. Similar to the within-network connections described above, tracts connecting the lOTC to the PCS were consistently superior to tracts connecting the lOTC to the IFG (**Fig 4g**). This superior to inferior organization mirrors the spatial layout of neighboring math and reading fROIs on the cortical surface (**Fig. 1b**).

**Figure 4.**
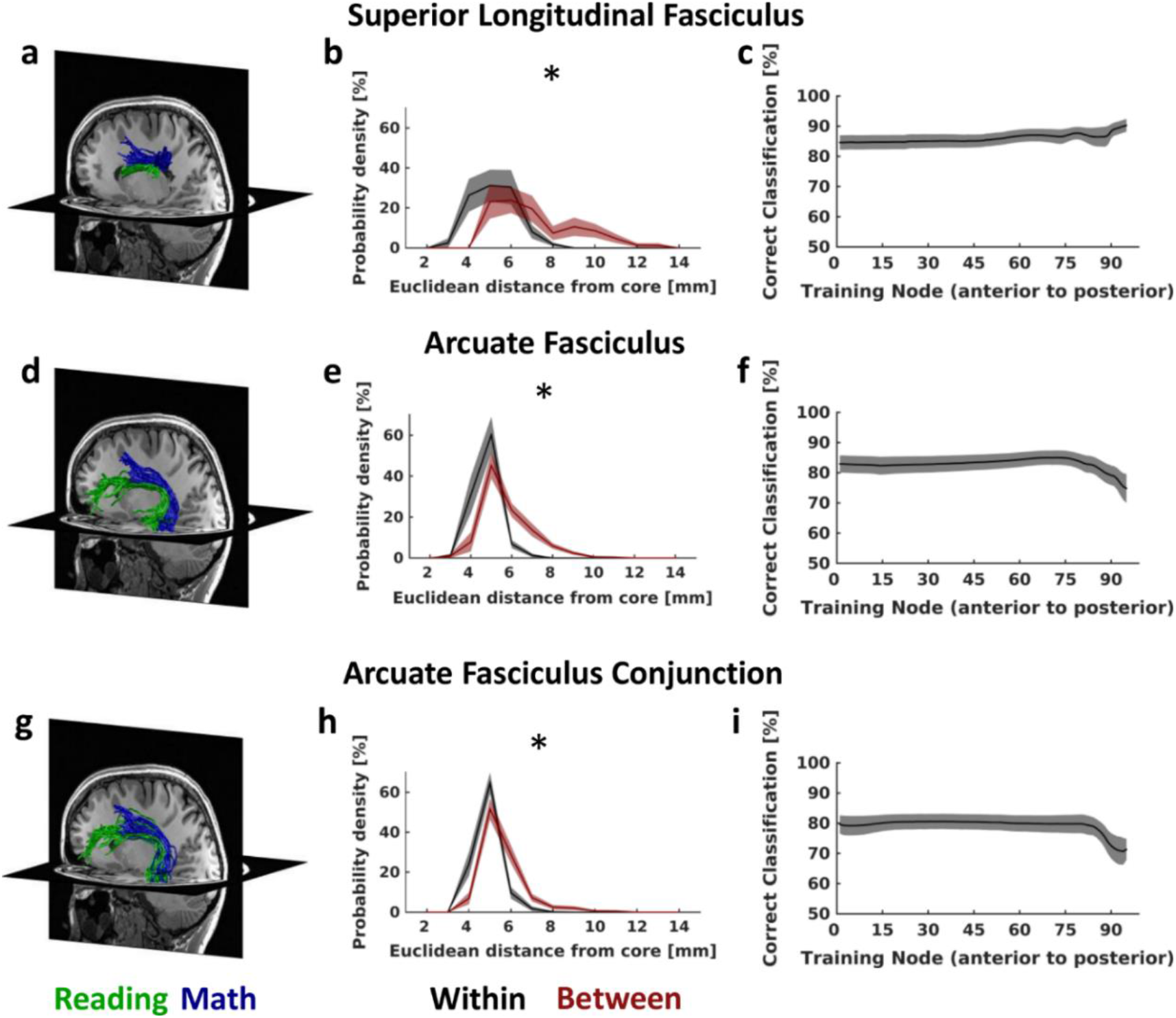
Pairwise connections of the reading and the math networks are segregated and parallel in the SLF and the AF. **(a-c)**: SLF tracts connecting IFG and SMGr in the reading network and PCS and SMGm in the math network. **(d-f)**: AF tracts connecting the IFG and STS in the reading network and the PCS and ITG in the math network. **(g-h)**: AF tracts connecting the lOTC fROI identified in the conjunction analysis with both the IFG in the reading network and the PCS in the math network. **(a,d,g)**: Math (blue) and reading (green) tracts of the SLF and AF in a representative individual subject showing the spatial segregation of these tracts. **(b,e,h)**: Euclidean distance in mm (derived from x,y,z coordinates) of all tracts relative to the core (mean) tract, within-network (black) and between-network (maroon). The distance was calculated across all tracts; the plot shows the mean across all nodes ±SEM. * Distributions differ significantly, p<0.05. **(c,f,i)**: Performance of a linear SVM classifying math and reading tracts within the SLF and AF based on their spatial location. Data show mean classification accuracy across nodes ±SEM. *Abbreviations:* IFG=inferior frontal gyrus, PCS=precentral sulcus, SMGr=reading fROI in supramarginal gyrus, SMGm=math fROI in supramarginal gyrus, STS=superior temporal sulcus, ITG=inferior temporal gyrus, lOTC=lateral occipito-temporal cortex, AF=arcuate fasciculus, SLF=superior longitudinal fasciculus.

To validate and quantify this segregation, we sectioned in each subject the SLF and AF to 100 equal-sized bins, referred to as nodes, and conducted two additional analyses:

(1) We measured the distribution of distances (Euclidean distance in mm) between individual tracts from the core tract (mean tract) of the network they belong to vs. the core tract of the other network. We reasoned that if tracts of the math and reading networks are segregated, distances to the within-network core tract should be smaller than to the between-network core tract. In contrast, if the tracts are intertwined, these distances should not be significantly different. Results show that individual tracts are significantly closer to the within-network core tract, compared to the core tract of the other network in both the SLF (two-sample Kolmogorov-Smirnov test on distance distributions: p<0.0001; paired t-test on mean distance across tract: p=0.0006, t(17)=4.22) and the AF (two-sample Kolmogorov-Smirnov test on distance distributions: p<0.0001; paired t-test on mean distance across tract: p<0.0001, t(17)= 6.12) (**Fig. 4b,e**).

In the AF, tracts connecting the conjunction fROI in the lOTC with the IFG (reading network) and the PCS (math network) were also segregated (**Fig. 4h**, two-sample Kolmogorov-Smirnov test on distance distributions: p<0.0001; paired t-test on mean distance across tract: p<0.0001, t(16)= 8.46).

(2) We used an independent classifier approach to evaluate if reading- and math-related fWMTs are spatially segregated at the entire length of the fascicles or only in a restricted region. We reasoned that if they are segregated at each node, a classifier should be able to determine if tracts belong to either the math or reading network solely based on their spatial location within the fascicle. To test this prediction, at each node, we trained a linear support vector machine (SVM) to distinguish math tracts from reading tracts based on their location (training: x,y,z coordinates of all tracts at a node). Then, we tested how well the SVM classifies new data based on tract coordinates from a more posterior node (we chose the fifth more posterior node relative to the training node, to ensure independence of training and test data). Across subjects and nodes, classification of tracts as belonging to either the reading or the math network was greater than 80% correct in the SLF and greater than 70% in the AF (**Fig. 4c,f**). In the AF decoding accuracy dropped towards the posterior portion of the tract (**Fig. 4f**). Notably, the average classification across both fascicles was significantly higher than the 50% chance level (SLF: p<0.0001, t(17) = 17.47; AF: p<0.0001, t(17) = 13.28).

When evaluating tracts within the AF that connect the conjunction fROI in the lOTC with the IFG (reading network) and the PCS (math network) (**Fig. 4i**), decoding accuracy again dropped towards the posterior end of the tract. Decoding accuracy still remained greater than 65%, and was significantly higher than the 50% chance level (p<0.0001, t(16) = 10.63).

Similar results were also obtained for the right hemisphere (**Supplementary Fig. 17**), for constant-size spherical fROIs (**Supplementary Fig. 18**) and when pairwise tracts were identified by using one fROI as a seed and the other fROI as a target for tractography (**Supplementary Fig. 19**). Overall, our analyses show that tracts associated with math and reading remain segregated, and largely parallel to each other within the SLF and the AF. Within both fascicles, tracts of the math network are located superior to tracts of the reading network.

### Reading tracts show faster T_1_ than math tracts

We next asked if there are structural differences between reading- and math-related fWMT within the SLF and AF. To address this question, we used qMRI to determine T_1_ of the reading and math fWMTs. T_1_ is inversely correlated with myelination (lower T_1_ is associated with higher myelin content). We also evaluated macromolecular tissue volume fraction (MTV), which indicate the fraction of non-water tissue in each voxel (**Supplementary Fig. 20**).

We first measured the average T_1_ of math- and reading-related tracts across the length of the fascicles. In the SLF (**Fig. 5a**) and in the AF (**Fig. 5d**) the average T_1_ of reading-related tracts was significantly lower compared to math-related tracts (SLF: p<0.0001, t(17)=5.72; AF: p=0.02, t(17)=2.66). Since the SLF and AF are long fascicles, we also tested for local differences across the tracts. For this, we segmented each tract to 100 nodes in each subject and then measured T_1_ along the tracts. Examination of the distributions of T_1_ values across the nodes of these tracts showed lower T_1_ in the fMWTs of the reading network compared to the math network in both the SLF (two-sample Kolmogorov-Smirnov test: p<0.0001, **Fig. 5b**) and the AF (two-sample Kolmogorov-Smirnov test: p<0.001, **Fig. 5e**). Examination of the spatial distribution of T_1_ values revealed differences across the length of the fascicles. In both the SLF and the AF, T_1_ differences were more pronounced towards the anterior end of the fascicle, compared to its posterior end (**Fig. 5c,f**).

In addition to the within-network connections described above, we also evaluated, within the AF, T_1_ of tracts connecting the conjunction fROI in the lOTC with either the IFG (reading network) or the PCS (math network) (**Fig. 5i**). We found a slight but significant differences in the distributions of T_1_ of math- and reading-related tracts (**Fig. 5h**, two-sample Kolmogorov-Smirnov test: p=0.008), but there was no significant difference in the mean T_1_ of these tracts (**Fig. 5g**, p=0.13, t(16) = 1.58).

Finally, a similar pattern of results was observed when evaluating MTV (**Supplementary Fig. 20**), in the right hemisphere (**Supplementary Fig. 21**), as well as for control analyses using constant-size spherical fROIs (**Supplementary Fig. 22**) and direct fROI to fROI tractography (**Supplementary Fig. 23**). The shorter T_1_ found along the SLF and AF for tracts associated with reading suggests that these tracts are more heavily myelinated then those associated with math.

**Figure 5.**
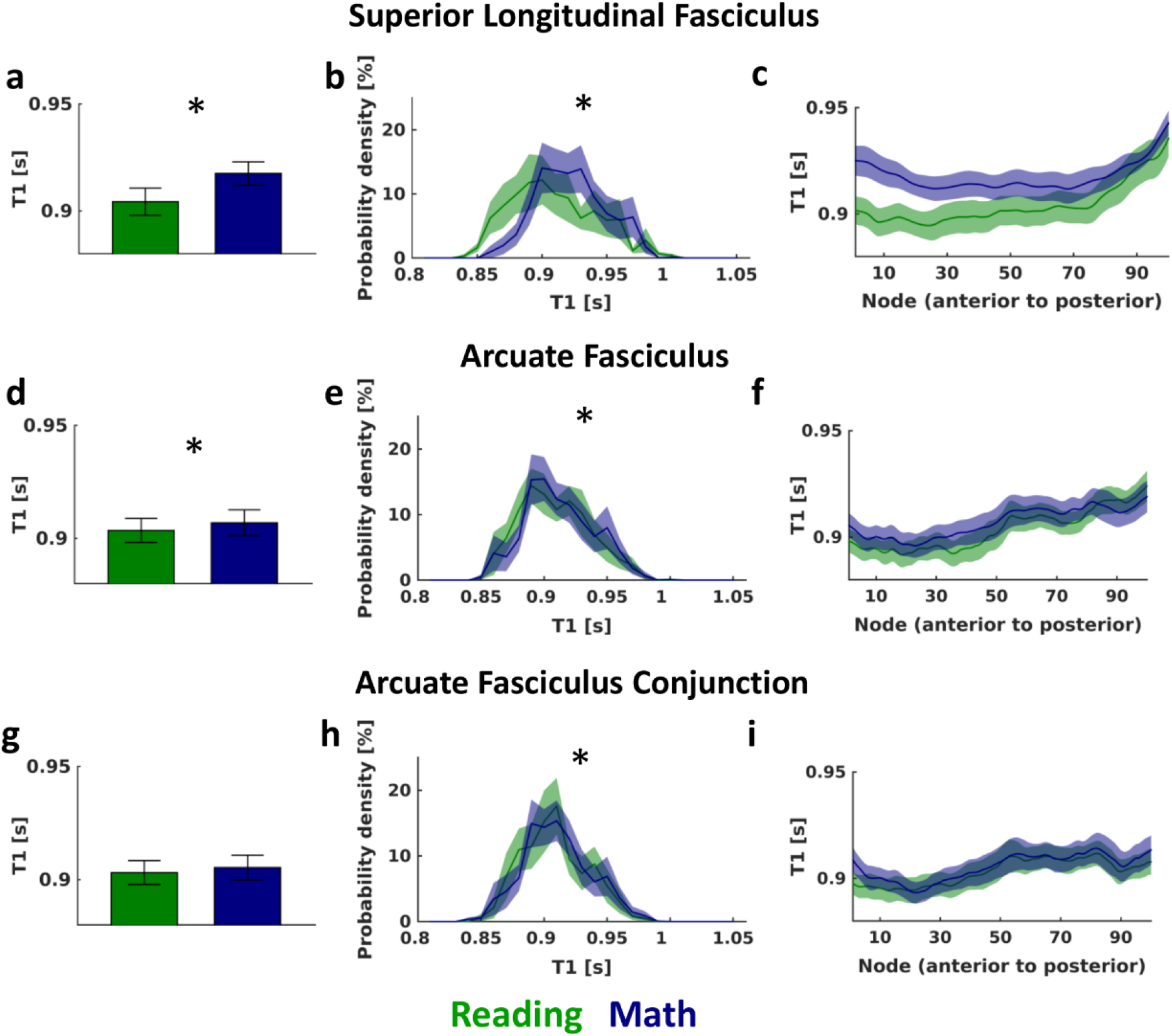
Tracts associated with reading show faster proton relaxation time (T_1_) than those associated with math in the SLF and the AF. **(a-c)**: T_1_ measurements for SLF tracts connecting IFG and SMGr in the reading network (green) and PCS and SMGm in the math network (blue). **(d-f)**: T_1_ measurements for AF tracts connecting the IFG and STS in the reading network (green) and the PCS and ITG in the math network (blue). **(g-i)**: T_1_ measurements for AF tracts connecting the lOTC conjunction fROI with the IFG in the reading network (green) and the PCS in the math network (blue). Left **(a,d,g)**: Average T_1_ for reading- and math-related tracts in the SLF and the AF. Bar graph shows mean across subjects ± SEM. *: T_1_ for math and reading related tracts differs significantly, p<0.05. Middle **(b,e,h)**: Distribution of T_1_ values across all tracts. Distributions were calculated within each subject and node; the plot shows the mean across all nodes ± SEM. *: Distributions differ significantly, p<0.05. Right **(c,f,i)**: Average T_1_ for reading- and math-related tracts along the SLF and the AF. Line graph shows mean across subjects ± SEM. *Abbreviations:* IFG=inferior frontal gyrus, PCS=precentral sulcus, SMGr=reading fROI in supramarginal gyrus, SMGm=math fROI in supramarginal gyrus, STS=superior temporal sulcus, ITG=inferior temporal gyrus, lOTC=lateral occipito-temporal cortex, SLF=superior longitudinal fasciculus, AF=arcuate fasciculus.

## Discussion

In the current study, we addressed a fundamental gap in knowledge in human brain function: what are the shared and dissociated gray and white matter substrates of math and reading, the two most essential skills every child is expected to acquire in school. Our multimodal data revealed three key findings. First, neighboring gray matter regions in the math and reading networks show similar white matter connectivity. Second, the AF and SLF are important fascicles for within-network connectivity in both the reading and the math network. Third, within the SLF and AF, tracts associated with math and reading are segregated, and show significant structural differences. Our data thereby opens a new avenue of research focused on understanding how sub-bundles within fascicles may contribute to human behavior.

Several methodological innovations were key to the present study. First, in contrast to many prior studies which commonly used tensor-based tracking methods (except for^12,50,7^), we used spherical deconvolution. This allowed us to resolve white matter tracts close to the gray matter. Second, by combining fMRI and dMRI, and intersecting each subjects’ tracts with the GWMI directly underneath our fROIs, we were able to define the fWMT of the math and the reading networks within individual subjects. This produced a precise and comprehensive understanding of the white matter associated with multiple functional regions of the math and reading networks. In comparison, prior studies have either examined white matter^6,7^ or functional gray matter regions^6,27^ of math and reading in isolation, evaluated white and gray matter substrates in distinct groups of participants^35^, or have examined the white matter connections of a single region in the reading network, the VWFA^18–20^. Third, the present study is the first to apply qMRI to elucidate structural properties of fascicles involved in math and reading. Prior investigations of white matter tracts of math and reading reported FA, mean diffusivity, or directional diffusivity of these tracts (e.g. ^8–11,13,15^). While these metrics provide valuable characterization of white matter and its development^45^, the relation between these metrics and the underlying microstructure is complex and not well understood^51,52^. In contrast T_1_ is correlated with myelin content^44^, thereby providing important insights about a fundamental microstructural component of these tracts. Fifth, by designing an experiment that uses identical stimuli for three different tasks, which are matched in their working memory load and the amount of verbalization they elicit, we were able to distil cortical regions that are involved in math, reading or both, while controlling for stimulus differences as well as general cognitive demands.

It should be noted though, that the fascicles of the math and reading networks reported here likely do not contain the entire white matter of these networks, for three reasons. First, in addition to the within-network connections described here, the cortical regions activated during reading and math likely also connect to regions outside their respective networks. In fact, our data does indeed show several significant connections between the math and reading networks (**Fig. 3**), although, on average, we found stronger within-than between-network connectivity. Second, there are likely additional white matter tracts associated with each region beyond long-range fascicles (e.g. shorter tracts, U-fibers), and these have not been considered here. Third, here we focused on addition, and did not investigate neural substrates of other arithmetic operations (subtraction, multiplication, and division) or broader mathematical abilities. Previous work has shown that different arithmetic operations vary in their neural substrates^25,53,54^ and that addition of small numbers relies more heavily on arithmetic fact retrieval compared to other arithmetic operations such as subtraction^55^. Arithmetic fact retrieval, in turn, has been linked to phonological processing^25,56,57^, which is also involved in reading (for review see^6,58^). These findings are in line with the observation that, compared to other arithmetic tasks, brain activations induced by addition show more overlap with brain activations induced by reading^2^. Here, we chose to investigate addition, using number-letter morphs, in order to match the mathematical and reading tasks as much as possible. Future studies that examine the gray and white matter substrates of other arithmetic operations and broader mathematical abilities will shed important light on which components of the revealed math network are specific to addition and which components extent beyond this arithmetic task.

Interestingly, by investigating both math and reading within the same participants, we found that cortical regions preferentially activated during these tasks are often neighboring (**Fig. 1b**). This neighboring relationship was observed in (i) the frontal cortex (reading centered on IFG and math on PCS), (ii) the lateral occipito-temporal cortex (reading centered on OTS/STS and math on ITG), and (iii) the SMG. Interestingly, gray matter regions involved in math were found to generally be located superior to regions involved in reading, which mirrors the superior-to-inferior arrangement of tracts associated with math and reading, respectively, in the AF and SLF. Further, as prior research suggests that white matter development precedes and predicts the location of functional regions involved in reading, such as the VWFA,^59^ our data highlights the possibility that long range white matter tracts may also determine the locations of cortical regions involved in math. Future developmental research could test this hypothesis.

Our study yields novel insights on the fascicles of the reading and math networks. First, we show that the SLF and AF are shared across the math and reading networks. Second, even as the SLF and AF are key fascicles for both tasks, they each contain separate sub-bundles for reading or math. Specifically, analogous to separate lanes on a highway, parallel and segregated tracts within these fascicles are part of either the reading or the math network (**Fig. 4, Supplementary Fig. 14-16**). These distinct sub-bundles for math and reading within the SLF and AF suggest that the white matter connections of the math and reading networks are more spatially specific than was previously believed. Third, strikingly, we found structural differences within math and reading sub-bundles in the AF and SLF. That is, T_1_ was shorter for reading than math tracts.

These findings have two important implications. First, our data shows that, even though math and reading involve several shared cognitive processes, such as the encoding of visual stimuli and working memory^1^, they are processed largely in parallel. This, in turn, suggests that improvements in one skill may not translate to the other skill, unless this improvement is linked to broad changes that transcend entire fascicles. Second, the faster T_1_ of the reading tracts than the math tracts within the SLF and AF suggests more substantial myelination of the former than the latter (**Fig. 5**). Notably, as reading is practiced more frequently and intensely than math during childhood^60^, and myelination is dependent on neural activity^61^, our findings raise the intriguing possibility that the amount of learning and its resultant neural activity may affect myelination of specific white matter tracts within fascicles.

This last finding makes an interesting prediction for potential links between white matter properties and math and reading skills: We hypothesize that the properties of the inferior and superior sections of the SLF and AF may independently contribute to reading and math performance. That is, if myelination improves transmission of information across distributed networks, then T_1_ of the inferior portion of the SLF and AF may correlate with reading ability, while T_1_ of the superior portions of these fascicles may correlate with math ability. Accordingly, atypical myelination of these tracts within the superior and inferior portion of the SLF and AF during development may also be associated with math or reading learning disabilities, respectively. Future studies with clinical populations that simultaneously evaluate the neural substrates of both math and reading learning disabilities can test these hypotheses. Further, we predict that if neural activity promotes myelination, then people with intensive practice in one of these tasks, such as expert mathematicians (e.g.^62^), may show lower T_1_ in the respective tracts compared to lay people. These predictions will be particularly relevant for studies evaluating the efficacy of interventions aimed at improving math and reading skills (e.g.^29,63,64^).

Crucially, the structural differences within the SLF and AF observed in the current study also have implications beyond math and reading. While structural differences within large tracts have recently been shown in the optic radiation, where an anterior sub-bundle that includes Meyer’s loop has higher T_1_ than the entire optic radiation^65^, the present study is the first to reveal such structural differences between functionally-defined tracts within fascicles. Our data thus encourages a novel research direction that links quantitative properties of functionally-defined sub-bundles to human behavior. That is, we believe that understanding the relationship between white matter properties and brain function (not only in reading and math^8–10,23,5^, but in a broad range of functions including face processing^66^, working memory^67^, and attention^68^) may be improved if white matter is defined more precisely, by linking it to the specific cortical regions that support each function.

In conclusion, our data shows striking functional and structural segregation of mathematical processing and reading in the human brain. These findings have implications for our understanding of the neural underpinning of math and reading as well as the link between white matter properties and human behavior more broadly.

## Methods

### Participants

20 volunteers (10 female, mean age ±SE: 27±1 years, 1 left-handed) were recruited from Stanford University and surrounding areas and participated in two experimental sessions. Subjects gave their informed written consent and the Stanford Internal Review Board on Human Subjects Research approved all procedures.

#### Functional MRI of math and reading related regions

##### Data acquisition and preprocessing

###### Anatomical MRI

A whole-brain, anatomical volume was acquired, once for each participant, using a T1-weighted BRAVO pulse sequence (resolution: 1mm x 1 mm x 1 mm, TI=450 ms, flip angle: 12°, 1 NEX, FoV: 240 mm). The anatomical volume was segmented into gray and white matter using FreeSurfer (http://surfer.nmr.mgh.harvard.edu/). with manual corrections using ITKGray (http://web.stanford.edu/group/vista/cgi-bin/wiki/index.php/ItkGray). From this segmentation, each participant’s cortical surface was reconstructed. Each participant’s anatomical brain volume was used as the common reference space for all analyses, which were always performed in individual native space.

###### Functional MRI

fMRI data was collected at the Center for Cognitive and Neurobiological Imaging at Stanford University, using a GE 3 tesla Signa Scanner with a 32-channel head coil. We acquired 48 slices covering the entire cortex using a T2*-sensitive gradient echo sequence (resolution: 2.4 mm x 2.4 mm x 2.4 mm, TR: 1000 ms, TE: 30 ms, FoV: 192 mm, flip angle: 62°, multiplexing factor of 3). A subset (N=12) of the fMRI data were also used for our previous study^36^.

The functional data was analyzed using the mrVista toolbox (http://github.com/vistalab) for Matlab, as in previous work^36^. In short, the data was motion-corrected within and between scans and then manually aligned to the anatomical volume. The manual alignment was optimized using robust multiresolution alignment^69^. No smoothing was applied. The time course of each voxel was high-pass filtered with a 1/20 Hz cutoff and converted to percentage signal change. A design matrix of the experimental conditions was created and convolved with the hemodynamic response function (HRF) implemented in SPM (http://www.fil.ion.ucl.ac.uk/spm) to generate predictors for each experimental condition. Response coefficients (betas) were estimated for each voxel and each predictor using a general linear model (GLM).

##### Stimuli and design

In the fMRI experiment, we presented well-controlled character-like stimuli, which could be used for a reading task, a math task, and a color memory task (**Fig. 1a**). These stimuli allowed us to define both math- and reading-related brain regions within the same experiment, while keeping the visual input constant. Details on the experimental design can be found in a previous study^36^. In brief, at the beginning of each trial, subjects were presented with a cue (“Add”, “Read”, or “Color”), indicating which task they should perform. In the adding task, participants were asked to sum the values of the stimuli and to indicate the correct sum. In the reading task, subjects were instructed to read the word in their head, and to indicate which word had been presented. Finally, in the color task, participants were asked to memorize the color of the stimuli and to indicate which color was shown during the trial. After the cue, 4 images were shown sequentially, followed by an answer screen. Each image was a morph of a number and a letter. All images in a trial were either number morphs (N, >80% number + <20% letter) or letter morphs (L, >80% letter + <20% number), i.e. stimuli that mostly contained information from one category, but held just enough evidence from the other category to be recognizable as both letters and numbers. The same stimuli appeared in all tasks. The answer screen was presented for 2 seconds and showed the correct answer as well as one incorrect answer at counterbalanced locations left and right of fixation. Participants performed 6 runs, each lasting six minutes, and the task order was randomized across runs and participants. Prior to the experiment, subjects were given training to ensure that they could perform the task with at least 80% accuracy.

##### Functionally-defined gray matter regions

Reading- and math-related gray matter regions of interest (fROIs) were defined in each participant’s cortical surface using both functional and anatomical criteria. For example, for our IFG fROI we took only those voxels that (i) showed the relevant task preference beyond the threshold of T≥3 and (ii) fell within the inferior frontal gyrus. The resulting fROIs were labeled according to their anatomical location. Reading-related fROIs consist of voxels that showed higher responses in the reading than the math and the color task (T≥3, voxel level), while math-related fROIs contain voxels which showed higher responses in the math than the reading and the color task (T≥3, voxel level). We also identified regions that are involved in both math and reading using a conjunction analysis (math>color ∩ reading > color, T≥3, voxel level) and extracted the response profile of the resulting fROIs (**Supplementary Fig. 8**). For all analyses, we report data from regions that showed a reliable preference for a given task across subjects. That is, we report regions that could be identified in the left hemisphere in at least 90% of the participants. In other words, while in a given individual there may be additional voxels that respond preferentially during reading and/or during math, here we focus on the most consistent activations. Given that reading related fROIs were more commonly found in the left hemisphere, the main text focuses on this hemisphere, whereas right hemisphere data is presented in the Supplementary Material (**Supplementary Fig. 2, 4, 5, 7, 10, 12, 17, 21**).

In addition to math- and reading-related regions, we also defined fROIs involved in color memory, which were used to determine a chance level for pairwise connections in the math and reading networks. Color-preferring voxels were identified in the medial aspect of the fusiform gyrus and showed significantly higher responses during the color task than the other two tasks (T≥3, voxel level). These voxels were frequently divided into three distinct subregions (likely corresponding to color patches Ac, Cc, and Pc^70^). Given that these regions are proximal, here we took the union of these color patches (left hemisphere: N=19, size ±SE: 318±75 mm^3^; right hemisphere: N=18, size ±SE: 277±64 mm^3^).

#### Diffusion MRI of the math and reading networks

##### Data acquisition and processing

Diffusion-weighted MRI (dMRI) data was collected in the same participants during a different day than the fMRI data, at the same facility and with the same 32-channel head-coil. DMRI was acquired using a dual-spin echo sequence in 96 different directions, 8 non-diffusion-weighted (b=0) images were collected, 60 slices provided full head coverage (resolution: 2 mm × 2 mm × 2 mm, TR: 8000 ms, TE: 93.6 ms, FoV: 220 mm, flip angle: 90°, b: 2000 s mm^-2^).

DMRI data was preprocessed using a combination of tools from mrTrix3 (github.com/MRtrix3/mrtrix3) and mrDiffusion toolbox (http://github.com/vistalab) for Matlab. First, we denoised the data using i) a principal component analysis, ii) Rician based denoising, and iii) Gibbs ringing corrections^71–73^. Second, we corrected for eddy currents using FSL (https://fsl.fmrib.ox.ac.uk/) and we performed bias correction using ANTs^74^. Third, dMRI data was registered to the average of the non-diffusion weighted images and aligned to the corresponding high-resolution anatomical brain volume using rigid body transformation. Fourth, voxel-wise fiber orientation distributions (FOD) were calculated using constrained spherical deconvolution (CSD)^75^ with up to 8 spherical harmonics (lmax=8). The FODs were then used for tractography.

#### Tractography

Ensemble tractography was performed on the processed dMRI data and consisted of 3 main steps: We (1) created multiple connectomes that varied in their allowed angle, (2) concatenated these candidate connectomes into one large ensemble connectome, and (3) automatically labelled major fascicles in the ensemble connectome.

1. Candidate connectome generation: We used MRtrix3^39,76^ (RC3, http://www.mrtrix.org/) to generate 5 candidate connectomes which varied in the maximum angle (2.25°, 4.5°, 9°, 11.25°, 13.5°). The goal of this approach was to generate candidate connectomes with tracts with different degrees of curviness, rather than limiting the connectome to one particular set of parameters.^38^ For each connectome, we used probabilistic fiber tracking with the following parameters: algorithm: IFOD1, step size: 0.2 mm, minimum length: 4 mm, maximum length: 200 mm, FOD amplitude stopping criterion: 0.1. We used anatomically constrained tractography (ACT)^77^, which utilizes information of different tissue types from the FreeSurfer (https://surfer.nmr.mgh.harvard.edu/) segmentation of each participant’s high-resolution anatomical scan to optimize tractography. ACT also allowed us to identify the gray-white matter interface (GWMI) directly underneath the fROIs. Seeds for tractography were randomly placed within this interface. This enabled us to focus on those fiber tracts that reach the gray matter. Each candidate connectome consisted of 500,000 streamlines.
2. The 5 candidate connectomes were concatenated into one ensemble connectome containing a total of 2,500,000 streamlines using custom Matlab code available in the Fascicle Analysis Toolbox (FAT) (https://github.com/VPNL/fat).
3. We used Automated Fiber Quantification^40^ (AFQ, https://github.com/yeatmanlab/AFQ) to segment the ensemble connectome of each participant into 13 well-established major fascicles, most of them are bilateral (**Supplementary Fig. 9**). The resulting classified connectome was optimized by removing tracts that were located more than 4 standard deviations away from the mean of their respective fascicle (similar to e.g. ^40,45^). We conducted all subsequent analyses on these classified white matter tracts within the 13 major fascicles, as we were interested in identifying large-scale, whole-brain networks involved in reading and adding.

##### Functionally-defined white matter tracts (fWMT)

###### fWMT of each fROI

To identify the tracts associated with each math and reading fROI, we intersected the classified tracts within 13 major fascicles with the GWMI directly adjacent to each of the math- and reading-related fROIs. This yielded the functionally-defined white matter tracts (fWMT) of each fROI. We visualized the reading-(**Fig. 2a**) and math-(**Fig. 2b**) related fWMT, as well as those tracts that connect to an lOTC region identified in the conjunction analysis (**Fig. 2c**), in individual subjects. In all plots, tracts are color-coded by the fascicle they belong to; the plots are thresholded at a maximum of 50 tracts per fascicle to enable clearer visualization. We also quantified the distribution of the fWMT of each fROI across the 13 main fascicles (**Fig. 2a,b,c-middle**). Finally, we created a schematic representation of the fascicles each fROI connects to, where the width of the line represents the proportion of the total fWMT occupied by that fascicle (**Fig. 2a,b,c-right**).

###### Within-network connections

To identify the tracts associated with the math and reading networks, we identified pairwise connections between fROIs within each network. We intersected the fWMT of each fROI with the GWMI underneath each of the other fROIs of the same network (either math or reading) to identify tracts that connect to at least two fROIs of the network. For both reading (**Fig. 3a-d**) and math (**Fig. 3e-h**), we first visualized all pairwise fWMT in each individual subject, color coded by the fascicle they belong to (**Fig. 3-a,e** show a representative subject).

Next, we quantified the pairwise connections using the dice coefficient (DC^49^; **Fig. 3-b,f**):

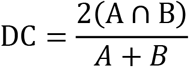

where A is all tracts that connect to one region, B is all tracts that connect to the second region, and A∩B is those tracts that connect to both regions.

The dice coefficient (DC) quantifies the similarity of two samples; a DC of 1 indicates complete overlap (i.e. each tract that connects to one region, also connects to the other region), while a DC of 0 indicates that there are no tracts that connect to both regions. We also estimated chance level DCs, by calculating the DC for pairwise fWMT between each of the fROIs of the math and the reading network to ventral regions activated during the color memory task (pairwise connections with the OTS were not included, due to its close anatomical proximity with fROIs involved in processing color). The average DCs of these pairwise connections were used as chance level DCs for each network, given that we expected connections to color-related regions to be irrelevant for participants’ math and reading skills. In addition, we also evaluated what percentage of the pairwise connections belong to each fascicle (**Fig. 3-c,g**). Finally, we created a schematic representation of these pairwise connections, where the width of the line is determined from the DC and the color of the lines indicate which fascicle contributed most strongly to this connection (**Fig. 3-d,h**).

###### Between-network connections

We also tested for between-network connections, using two approaches: (i) We identified and quantified the pairwise connections between the lOTC conjunction fROI and each non-neighboring fROIs in both the math and reading networks with the methods described above (**Fig. 3i-l**), and (ii) we identified connections between pairs of non-neighboring fROIs across networks using the same methods as above except that in each pair one fROI was from the reading network and the other fROI from the math network (**Fig. 3m-o**).

##### Quantification of segregation within fascicles

For those fascicles that contributed to significant pairwise connections in both the math and the reading network, we tested whether fWMT within these fascicles are segregated by network. Specifically, we evaluated if (i) within the SLF, tracts connecting SMGr to the IFG in the reading network are intertwined or segregated from tracts connecting SMGm to the PCS in the math network (**Fig. 4a-c**, tracts that connect to all 4 fROIs were excluded), (ii) within the AF, tracts connecting the STS and the IFG are intertwined or segregated from tracts connecting the ITG and the PCS (**Fig. 4d-f**, tracts that connect to all 4 fROIs were excluded), and (iii) within the AF, tracts connecting the lOTC fROI identified in the conjunction analysis with the IFG are intertwined or segregated from tracts connecting the same lOTC fROI with the PCS (**Fig. 4g-i**, tracts that connect to all 3 fROIs were excluded).

To do these analyses, we first visualized pairwise connections of the math and reading network within each fascicle, in each individual subject, to visually inspect their spatial layout (**Fig. 4-a,d,g and Supplementary Fig. 14-16**). Then, we resampled each tract in each subject to 100 equally spaced nodes (i.e. locations) between the way-point ROIs used by AFQ to define the fascicle. This procedure ensured that we have the same number of measurements per subject, even though the absolute length of the fascicles may vary across subjects. Finally, we quantified the segregation of fWMT of each network within each fascicle using two complimentary approaches: (1) we measured the distance of each tract from the core tract of within network-tracts, as well as, its distance from the core tract of the other network; then we tested whether the former is lower than the latter and (2) we used an independent classifier to test if, across nodes, fWMT can be identified as belonging to either the reading or the math network based on their anatomical location.

1. <u>Distance to core tract within and between networks</u>: We first calculated the core (mean) tract of the pairwise connections within the SLF and AF, separately for math and reading-related fWMT, using AFQ. Next, we measured, within each subject and at each node, how far away (Euclidian distance in mm, derived from x,y,z coordinates) each tract is from the core tract within its network and the core tract of the other network (**Fig. 4-b,e, h**). We expected tracts to be closer to the core tract of their own network if math- and reading-related fWMT are segregated within the fascicle, but equal distant from both core tracts if math- and reading-related fWMT are intertwined within the fascicle.
2. <u>Classification</u>: We also tested if, across the length of the fascicle, we can classify tracts as “math” or “reading” tracts based on their anatomical location. At each node and within each subject, the coordinates of all math and reading related tracts were used to train a linear support vector machine (SVM) classifier. The SVM from each node was used to classify tracts at the fifths more posterior node (we chose the fifths node rather than a neighboring node to ensure independence of training and test data) as either “math” or “reading” (**Fig. 4-c,f,i**). We expected the classifier to perform at chance (50% accuracy) if math and reading related tracts are intertwined, but significantly above chance if math and reading tracts are spatially segregated across the lengths of the fascicle.

#### Quantitative MRI of math and reading tracts

##### Data acquisition and preprocessing

Quantitative MRI (qMRI^41^) data was collected within the same session and with the same head coil as the dMRI data. T_1_ relaxation times were measured from four spoiled gradient echo images with flip angles of 4°, 10°, 20° and 30° (TR: 14 ms, TE: 2.4 ms). The resolution of these images was later resampled from 0.8×0.8×1.0 mm^3^ to 1mm isotropic voxels, and qMRI data was aligned with the high-resolution anatomical scan using rigid body transformation. We also collected four additional spin echo inversion recovery (SEIR) scans with an echo planar imaging read-out, a slab inversion pulse and spectral spatial fat suppression (TR: 3 s, resolution: 2 mm x 2 mm x 4 mm, 4 echo time set to minimum full, 2x acceleration, inversion times: 50, 400, 1200, and 2400 ms). The purpose of these SEIRs was to remove field inhomogeneities.

Both the spoiled gradient echo and the SEIR scans were processed using the mrQ software package (https://github.com/mezera/mrQ) for Matlab to estimate the proton relaxation time (T_1_) and macromolecular tissue volume (MTV) in each voxel, as in previous studies^41,78^. In brief, the mrQ analysis pipeline corrects for RF coil bias using the SEIRs scans, which produces accurate proton density (PD) and T_1_ fits across the brain. MrQ also produces maps of MTV, by calculating the fraction of a voxel that is non-water. T_1_ and MTV maps of each subject were co-registered to the same anatomical whole brain volume as dMRI and fMRI data.

##### Comparison of T_1_ for math and reading tracts

We used the T_1_ maps to evaluate tissue properties of tracts of the math or the reading network (**Fig. 5**; MTV data is presented in **Supplementary Fig. 20**). We focused on those fascicles that supported significant within-network connectivity in both networks. Within the SLF, we compared tracts connecting SMGr to the IFG in the reading network with tracts connecting SMGm to the PCS in the math network (**Fig. 5a-c**, tracts that connect to all 4 fROIs were excluded). Within the AF, we compared tracts connecting the STS and the IFG in the reading network with tracts connecting the ITG and the PCS in the math network (**Fig. 5d-f**, tracts that connect to all 4 fROIs were excluded). Within the AF, we also compared tracts connecting the lOTC conjunction fROI with the IFG in the reading network to those connecting the lOTC conjunction fROI to the PCS in the math network (**Fig. 5g-i**). We first evaluated the mean T_1_ values of each tract in each subject and tested if there are between-network differences (**Fig. 5-a,d,g**). Then, we resampled the tract to 100 equally-spaced nodes in-between the way-point ROIs used by AFQ to identify the fascicle. We evaluated the distribution of T_1_ values, using data from all nodes and subjects, and tested if the distribution varies between math and reading tracts (**Fig. 5-b,e,h**). Finally, we visualized the T_1_ of math and reading tracts across the different nodes, to determine if T_1_ differences are homogenous or heterogeneous across the length of the tract (**Fig. 5-c,f,i**).

<u>Control Analyses</u>: In addition to our main approach, we conducted 2 control analyses:

1. Constant-sized fROI. We replicated our analyses using constant size spherical ROIs (radius=7mm, this radius was chosen based on previous studies, e.g.^19,35^) which were centered on the fROIs, in order to ensure that differences in ROI size across participants and regions did not influence the identified math and reading networks. Results of these analyses are presented in the Supplementary Materials (**Supplementary Fig. 11, 13, 18, 22**).
2. Direct fROI-to-fROI tractography. We replicated the pairwise connections between fROIs analyzed in **Fig. 4–5** by tracking between the fROIs, in the sense that the GWMI underneath the anterior fROI in each pair was used as a seed and the GWMI of the other fROI was used as a target for tractography. The advantage of this approach is that it allows to better control the number of tracts seeded in the GWMI of each fROI. However, in contrast to our main approach, this control analysis only tracks between pairs of fROI and hence does not provide a picture of the entire math and reading networks. We used the following parameters for this control analysis: algorithm: IFOD1 with ACT, step size: 0.2 mm, minimum length: 4 mm, maximum length: 200 mm, FOD amplitude stopping criterion: 0.1, angle: 13.5°. We continued tracking between the fROIs until i) we found 100 tracts or ii) we attempted 100.000 times to seed tracts. Tracts were classified and cleaned using AFQ as described above.

##### Statistics

Repeated measures analyses of variance (ANOVAs) were used to test for accuracy and RT differences across tasks, and to test if fROIs identified in the conjunction analysis respond equally strongly to math and reading tasks. We used paired t-tests to evaluate if DCs, or decoding accuracies differed significantly from chance. We also used paired t-test to evaluate if there are T_1_ differences between math and reading related tracts. When more than one t-test was conducted, the statistical threshold was Bonferroni-adjusted to account for multiple comparisons. When comparing distributions, for example distribution of T_1_ values, we used the non-parametric two-sample Kolmogorov-Smirnov test to test for statistically significant differences.

##### Data availability

The data generated in this study will be made available upon reasonable request.

##### Code availability

The fMRI and qMRI data were analyzed using the open source mrVista software (available in GitHub: http://github.com/vistalab) and mrQ software (available in GitHub: https://github.com/mezera/mrQ) packages, respectively. The dMRI data were analyzed using open source software, including MRtrix3^39^ (http://www.mrtrix.org/) and AFQ^40^ (https://github.com/yeatmanlab/AFQ). We make the entire pipeline freely available; custom code for preprocessing, tractography and further analyses are available in github (github.com/scitran-apps/mrtrix3preproc; https://github.com/VPNL/fat). Code for reproducing all figures and statistics are made available in github as well (https://github.com/VPNL/mrLanes).

## Acknowledgements

This research was supported by the National Institute of Health (NIH; 1R01EY02391501A1), by the Deutsche Forschungsgemeinschaft (DFG; GR 4850/1-1) and by an Innovation Grant from the Stanford Center for Cognitive and Neurobiological Imaging (CNI). The authors would like to thank Brianna Jeska for her help with the data collection.

## Author Contribution

MG collected the data. ZZ developed the FAT toolbox used for diffusion and quantitative data analyses. MG and GLU analyzed the data. MG and KGS wrote the manuscript.

## Materials and Correspondence

Correspondence and material requests should be directed to Mareike Grotheer.

## References

1. Ashkenazi, S., Rubinsten, O. & De Smedt, B. Editorial: Associations between reading and mathematics: Genetic, brain imaging, cognitive and educational perspectives. Front. Psychol. 8, 600 (2017).

2. Evans, T. M., Flowers, D. L., Luetje, M. M., Napoliello, E. & Eden, G. F. Functional neuroanatomy of arithmetic and word reading and its relationship to age. Neuroimage 143, 304–315 (2016).

3. Mann Koepke, K. & Miller, B. At the Intersection of Math and Reading Disabilities: Introduction to the Special Issue. J. Learn. Disabil. 46, 483–489 (2013).

4. Wandell, B. A. & Le, R. K. Diagnosing the Neural Circuitry of Reading. Neuron 96, 298–311 (2017).

5. Shaywitz, S. E. & Shaywitz, B. A. Paying attention to reading: The neurobiology of reading and dyslexia. Dev. psychopathol. 20, 1329–1349 (2008).

6. Peters, L. & De Smedt, B. Arithmetic in the developing brain: A review of brain imaging studies. Dev. Cogn. Neurosci. 30, 265–279 (2018).

7. Matejko, A. A. & Ansari, D. Drawing connections between white matter and numerical and mathematical cognition: A literature review. Neurosci. Biobehav. Rev. 48, 35–52 (2015).

8. Yeatman, J. D. et al. Anatomical properties of the arcuate fasciculus predict phonological and reading skills in children. J. Cogn. Neurosci. 23, 3304–3317 (2011).

9. Vandermosten, M. et al. A tractography study in dyslexia: Neuroanatomic correlates of orthographic, phonological and speech processing. Brain 135, 935–948 (2012).

10. Su, M. et al. Alterations in white matter pathways underlying phonological and morphological processing in Chinese developmental dyslexia. Dev. Cogn. Neurosci. 31, 11–19 (2018).

11. Vanderauwera, J., Wouters, J., Vandermosten, M. & Ghesquière, P. Early dynamics of white matter deficits in children developing dyslexia. Dev. Cogn. Neurosci. 27, 69–77 (2017).

12. Zhao, J., Thiebaut de Schotten, M., Altarelli, I., Dubois, J. & Ramus, F. Altered hemispheric lateralization of white matter pathways in developmental dyslexia: Evidence from spherical deconvolution tractography. Cortex 76, 51–62 (2016).

13. Vanderauwera, J. et al. Neural organization of ventral white matter tracts parallels the initial steps of reading development: A DTI tractography study. Brain Lang. 183, 32–40 (2018).

14. Epelbaum, S. et al. Pure alexia as a disconnection syndrome: New diffusion imaging evidence for an old concept. Cortex 44, 962–974 (2008).

15. Yeatman, J. D., Dougherty, R. F., Ben-Shachar, M. & Wandell, B. A. Development of white matter and reading skills. Proc. Natl. Acad. Sci. 109, E3045–E3053 (2012).

16. Yeatman, J. D. et al. The vertical occipital fasciculus: A century of controversy resolved by in vivo measurements. Proc. Natl. Acad. Sci. 111, E5214–E5223 (2014).

17. Kay, K. N. & Yeatman, J. D. Bottom-up and top-down computations in word- and face-selective cortex. Elife 6, (2017).

18. Bouhali, F. et al. Anatomical Connections of the Visual Word Form Area. J. Neurosci. 34, 15402–15414 (2014).

19. Yeatman, J. D., Rauschecker, A. M. & Wandell, B. A. Anatomy of the visual word form area: Adjacent cortical circuits and long-range white matter connections. Brain Lang. 125, 146–155 (2013).

20. Lerma-Usabiaga, G., Carreiras, M. & Paz-Alonso, P. M. Converging evidence for functional and structural segregation within the left ventral occipitotemporal cortex in reading. Proc. Natl. Acad. Sci. 115, 201803003 (2018).

21. Dehaene, S. & Cohen, L. The unique role of the visual word form area in reading. Trends in Cognitive Sciences 15, 254–262 (2011).

22. Cohen, L. et al. The visual word form area. Brain 123, 291–307 (2000).

23. Gaillard, R. et al. Direct Intracranial, fMRI, and Lesion Evidence for the Causal Role of Left Inferotemporal Cortex in Reading. Neuron 50, 191–204 (2006).

24. Harvey, B. M., Klein, B. P., Petridou, N. & Dumoulin, S. O. Topographic representation of numerosity in the human parietal cortex. Science (80-.). 341, 1123–1126 (2013).

25. Van Beek, L., Ghesquière, P., Lagae, L. & De Smedt, B. Left fronto-parietal white matter correlates with individual differences in children’s ability to solve additions and multiplications: A tractography study. Neuroimage 90, 117–127 (2014).

26. Piazza, M., Izard, V., Pinel, P., Le Bihan, D. & Dehaene, S. Tuning curves for approximate numerosity in the human intraparietal sulcus. Neuron 44, 547–555 (2004).

27. Arsalidou, M. & Taylor, M. J. Is 2+2=4? Meta-analyses of brain areas needed for numbers and calculations. Neuroimage 54, 2382–2393 (2011).

28. Tsang, J. M., Dougherty, R. F., Deutsch, G. K., Wandell, B. A. & Ben-Shachar, M. Frontoparietal white matter diffusion properties predict mental arithmetic skills in children. Proc. Natl. Acad. Sci. 106, 22546–22551 (2009).

29. Huber, E., Donnelly, P. M., Rokem, A. & Yeatman, J. D. Rapid and widespread white matter plasticity during an intensive reading intervention. Nat. Commun. 9, 2260 (2018).

30. Ben-Shachar, M., Dougherty, R. F. & Wandell, B. A. White matter pathways in reading. Curr. Opin. Neurobiol. 17, 258–270 (2007).

31. Klingberg, T. et al. Microstructure of temporo-parietal white matter as a basis for reading ability: Evidence from diffusion tensor magnetic resonance imaging. Neuron 25, 493–500 (2000).

32. Niogi, S. N. & McCandliss, B. D. Left lateralized white matter microstructure accounts for individual differences in reading ability and disability. Neuropsychologia 44, 2178–2188 (2006).

33. Beaulieu, C. et al. Imaging brain connectivity in children with diverse reading ability. Neuroimage 25, 1266–1271 (2005).

34. Li, Y., Hu, Y., Wang, Y., Weng, J. & Chen, F. Individual structural differences in left inferior parietal area are associated with schoolchildrens’ arithmetic scores. Front. Hum. Neurosci. 7, 844 (2013).

35. Klein, E., Moeller, K., Glauche, V., Weiller, C. & Willmes, K. Processing Pathways in Mental Arithmetic-Evidence from Probabilistic Fiber Tracking. PLoS One 8, 55455 (2013).

36. Grotheer, M., Jeska, B. & Grill-Spector, K. A preference for mathematical processing outweighs the selectivity for Arabic numbers in the inferior temporal gyrus. Neuroimage 175, 188–200 (2018).

37. Pestilli, F., Yeatman, J. D., Rokem, A., Kay, K. N. & Wandell, B. A. Evaluation and statistical inference for human connectomes. Nat. Methods 11, 1058–1063 (2014).

38. Takemura, H., Caiafa, C. F., Wandell, B. A. & Pestilli, F. Ensemble Tractography. PLoS Comput. Biol. 12, e1004692 (2016).

39. Tournier, J. D., Calamante, F. & Connelly, A. MRtrix: Diffusion tractography in crossing fiber regions. Int. J. Imaging Syst. Technol. 22, 53–66 (2012).

40. Yeatman, J. D., Dougherty, R. F., Myall, N. J., Wandell, B. A. & Feldman, H. M. Tract Profiles of White Matter Properties: Automating Fiber-Tract Quantification. PLoS One 7, (2012).

41. Mezer, A. et al. Quantifying the local tissue volume and composition in individual brains with magnetic resonance imaging. Nat. Med. 19, 1667–1672 (2013).

42. Lutti, A., Dick, F., Sereno, M. I. & Weiskopf, N. Using high-resolution quantitative mapping of R1 as an index of cortical myelination. Neuroimage 93, 176–188 (2014).

43. Sereno, M. I., Lutti, A., Weiskopf, N. & Dick, F. Mapping the Human Cortical Surface by Combining Quantitative T1 with Retinotopy†. Cereb. Cortex 23, 2261–2268 (2013).

44. Stüber, C. et al. Myelin and iron concentration in the human brain: A quantitative study of MRI contrast. Neuroimage 93, 95–106 (2014).

45. Yeatman, J. D., Wandell, B. A. & Mezer, A. A. Lifespan maturation and degeneration of human brain white matter. Nat. Commun. 5, 4932 (2014).

46. Weiner, K. S. & Grill-Spector, K. Neural representations of faces and limbs neighbor in human high-level visual cortex: Evidence for a new organization principle. Psychological Research 77, 74–97 (2013).

47. Daitch, A. L. et al. Mapping human temporal and parietal neuronal population activity and functional coupling during mathematical cognition. Proc. Natl. Acad. Sci. 113, E7277–E7286 (2016).

48. Baldauf, D. & Desimone, R. Neural mechanisms of object-based attention. Science (80-.). 344, 424–427 (2014).

49. Dice, L. R. Measures of the Amount of Ecologic Association Between Species. Ecology 26, 297–302 (1945).

50. Vanderauwera, J., Vandermosten, M., Dell’Acqua, F., Wouters, J. & Ghesquière, P. Disentangling the relation between left temporoparietal white matter and reading: A spherical deconvolution tractography study. Hum. Brain Mapp. 36, 3273–3287 (2015).

51. Mädler, B., Drabycz, S. A., Kolind, S. H., Whittall, K. P. & MacKay, A. L. Is diffusion anisotropy an accurate monitor of myelination?. Correlation of multicomponent T2relaxation and diffusion tensor anisotropy in human brain. Magn. Reson. Imaging 26, 874–888 (2008).

52. Jones, D. K., Knösche, T. R. & Turner, R. White matter integrity, fiber count, and other fallacies: The do’s and don’ts of diffusion MRI. NeuroImage 73, 239–254 (2013).

53. Rosenberg-Lee, M., Chang, T. T., Young, C. B., Wu, S. & Menon, V. Functional dissociations between four basic arithmetic operations in the human posterior parietal cortex: A cytoarchitectonic mapping study. Neuropsychologia 49, 2592–2608 (2011).

54. De Smedt, B., Holloway, I. D. & Ansari, D. Effects of problem size and arithmetic operation on brain activation during calculation in children with varying levels of arithmetical fluency. Neuroimage 57, 771–781 (2011).

55. Barrouillet, P., Mignon, M. & Thevenot, C. Strategies in subtraction problem solving in children. J. Exp. Child Psychol. 99, 233–251 (2008).

56. De Smedt, B., Taylor, J., Archibald, L. & Ansari, D. How is phonological processing related to individual differences in children’s arithmetic skills? Dev. Sci. 13, 508–520 (2010).

57. Hecht, S. A., Torgesen, J. K., Wagner, R. K. & Rashotte, C. A. The relations between phonological processing abilities and emerging individual differences in mathematical computation skills: A longitudinal study from second to fifth grades. J. Exp. Child Psychol. 79, 192–227 (2001).

58. De Smedt, B. Individual Differences in Arithmetic Fact Retrieval. Dev. Math. Cogn. 219–243 (2016). doi:10.1016/B978-0-12-801871-2.00009-5

59. Saygin, Z. M. et al. Connectivity precedes function in the development of the visual word form area. Nat. Neurosci. 19, 1250–1255 (2016).

60. Stacy, S. T., Cartwright, M., Arwood, Z., Canfield, J. P. & Kloos, H. Addressing the math-practice gap in elementary school: Are tablets a feasible tool for informal math practice? Front. Psychol. 8, 179 (2017).

61. Zatorre, R. J., Fields, R. D. & Johansen-Berg, H. Plasticity in gray and white: Neuroimaging changes in brain structure during learning. Nat. Neurosci. 15, 528–536 (2012).

62. Amalric, M. & Dehaene, S. Origins of the brain networks for advanced mathematics in expert mathematicians. Proc. Natl. Acad. Sci. 113, 4909–4917 (2016).

63. Keller, T. A. & Just, M. A. Altering Cortical Connectivity: Remediation-Induced Changes in the White Matter of Poor Readers. Neuron 64, 624–631 (2009).

64. Jolles, D. et al. Plasticity of left perisylvian white-matter tracts is associated with individual differences in math learning. Brain Struct. Funct. 221, 1337–1351 (2016).

65. Schurr, R. et al. Tractography optimization using quantitative T1 mapping in the human optic radiation. Neuroimage 181, 645–658 (2018).

66. Thomas, C. et al. Reduced structural connectivity in ventral visual cortex in congenital prosopagnosia. Nat. Neurosci. 12, 29–31 (2009).

67. Krogsrud, S. K. et al. Development of white matter microstructure in relation to verbal and visuospatial working memory—A longitudinal study. PLoS One 13, e0195540 (2018).

68. Klarborg, B. et al. Sustained attention is associated with right superior longitudinal fasciculus and superior parietal white matter microstructure in children. Hum. Brain Mapp. 34, 3216–3232 (2013).

69. Nestares, O. & Heeger, D. J. Robust multiresolution alignment of MRI brain volumes. Magn. Reson. Med. 43, 705–715 (2000).

70. Lafer-Sousa, R., Conway, B. R. & Kanwisher, N. G. Color-Biased Regions of the Ventral Visual Pathway Lie between Face- and Place-Selective Regions in Humans, as in Macaques. J. Neurosci. 36, 1682–1697 (2016).

71. Kellner, E., Dhital, B., Kiselev, V. G. & Reisert, M. Gibbs-ringing artifact removal based on local subvoxel-shifts. Magn. Reson. Med. 76, 1574–1581 (2016).

72. Veraart, J. et al. Denoising of diffusion MRI using random matrix theory. Neuroimage 142, 394–406 (2016).

73. Veraart, J., Fieremans, E. & Novikov, D. S. Diffusion MRI noise mapping using random matrix theory. Magn. Reson. Med. 76, 1582–1593 (2016).

74. Tustison, N. J. et al. N4ITK: Improved N3 bias correction. IEEE Trans. Med. Imaging 29, 1310–1320 (2010).

75. Tournier, J. D., Calamante, F. & Connelly, A. Robust determination of the fibre orientation distribution in diffusion MRI: Non-negativity constrained super-resolved spherical deconvolution. Neuroimage 35, 1459–1472 (2007).

76. Tournier, J.-D. et al. MRtrix3: A fast, flexible and open software framework for medical image processing and visualisation. bioRxiv 551739 (2019). doi:10.1101/551739

77. Smith, R. E., Tournier, J. D., Calamante, F. & Connelly, A. Anatomically-constrained tractography: Improved diffusion MRI streamlines tractography through effective use of anatomical information. Neuroimage 62, 1924–1938 (2012).

78. Gomez, J. et al. Microstructural proliferation in human cortex is coupled with the development of face processing. Science (80-.). 355, 68–71 (2017).

