## Supplemental Material for "Separate lanes for math and reading in the white matter highways of the human brain"

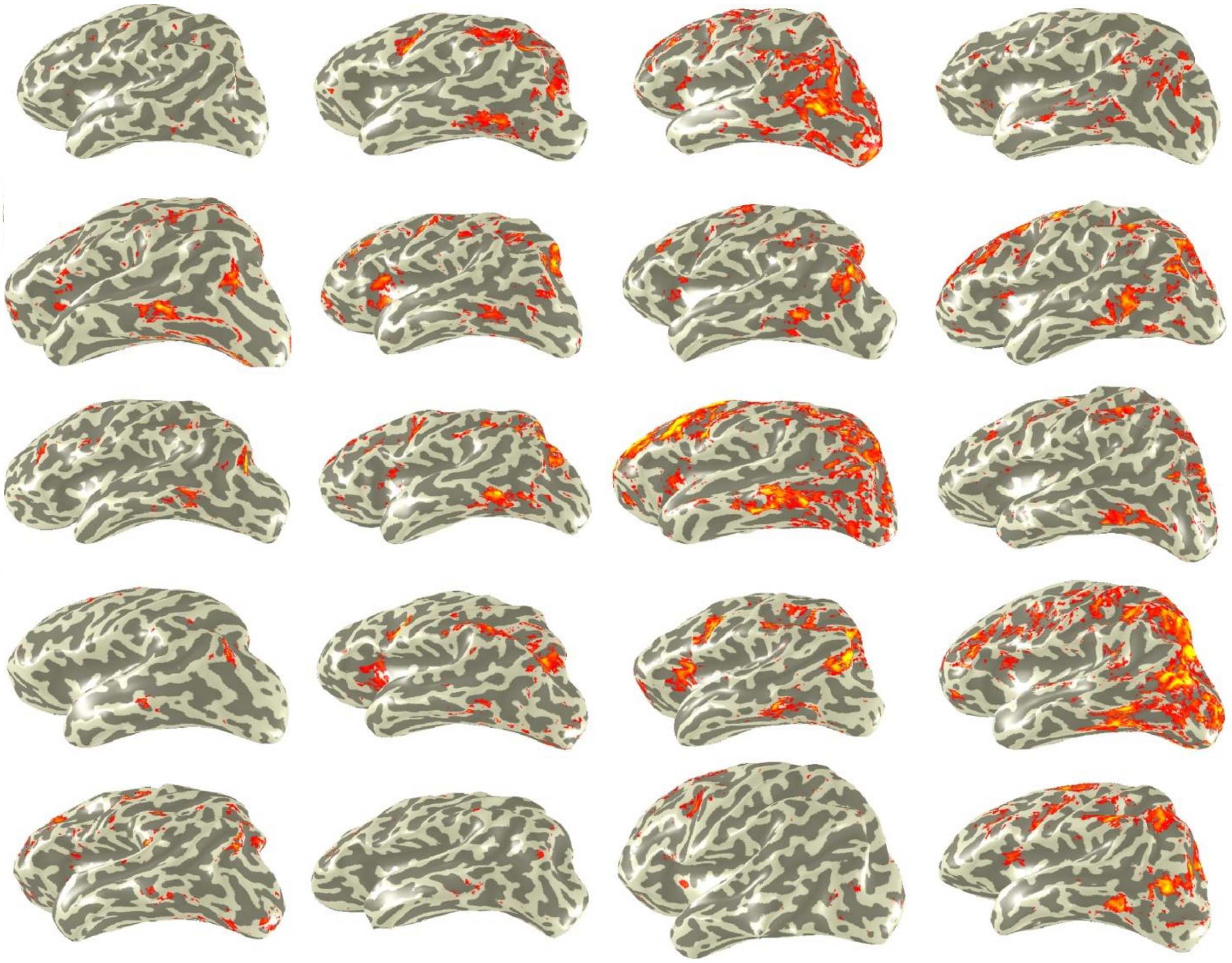

**Math > Reading + Color (T≥3)**

**Supplementary Fig. 1. Multiple regions in the brain show higher responses to the math task than the reading and color tasks.** BOLD responses contrasting the math task with the reading and the color tasks (T≥3) are presented in the left hemisphere of all subjects. Subjects are sorted by their response accuracy in the math task, from highest (*top left*) to lowest (*bottom right*) mean performance.

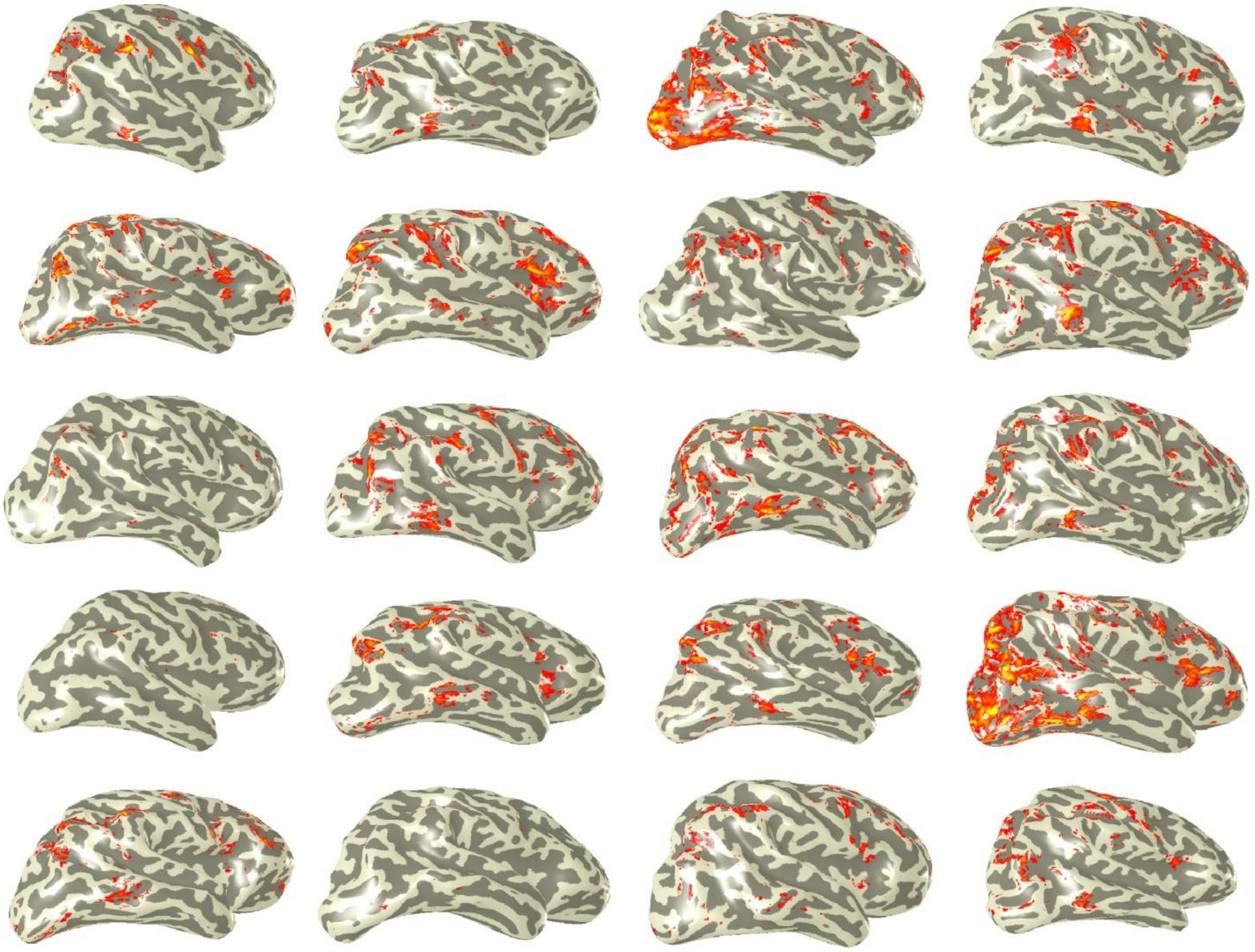

**Math > Reading + Color (T≥3)**

**Supplementary Fig. 2. Multiple regions in the brain show higher responses to the math task than the reading and color tasks.** BOLD responses contrasting the math task with the reading and the color tasks (T≥3) are presented in the right hemisphere of all subjects. Subjects are sorted by their response accuracy in the math task, from highest (*top left*) to lowest (*bottom right*) mean performance.

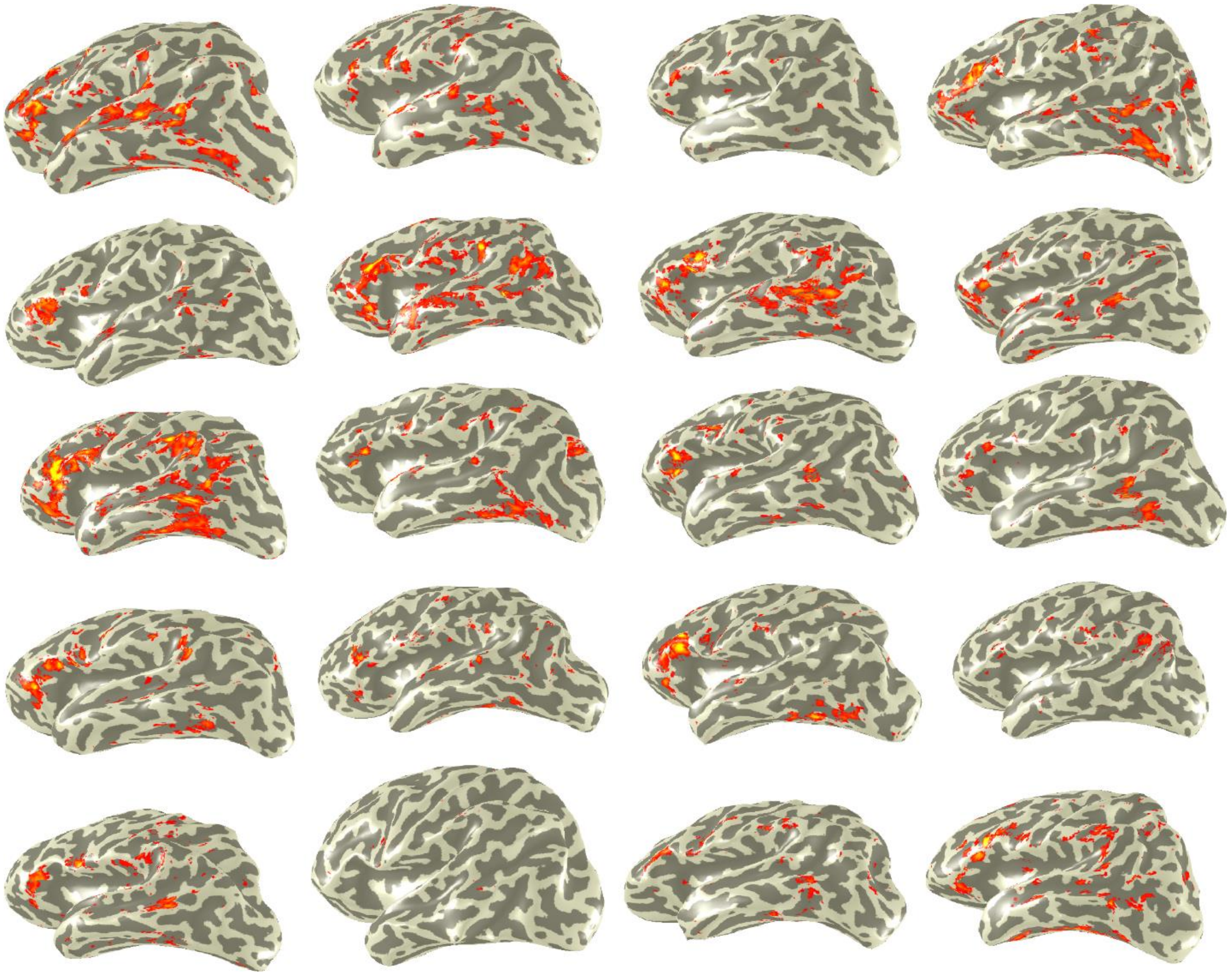

**Reading > Math+ Color (T≥3)**

**Supplementary Fig. 3. Multiple regions in the brain show higher responses to the reading task than the math and color tasks.** BOLD responses contrasting the reading task with the math and the color tasks (T≥3) are presented in the left hemisphere of all subjects. Subjects are sorted by their response accuracy in the math task, from highest (*top left*) to lowest (*bottom right*) mean performance.

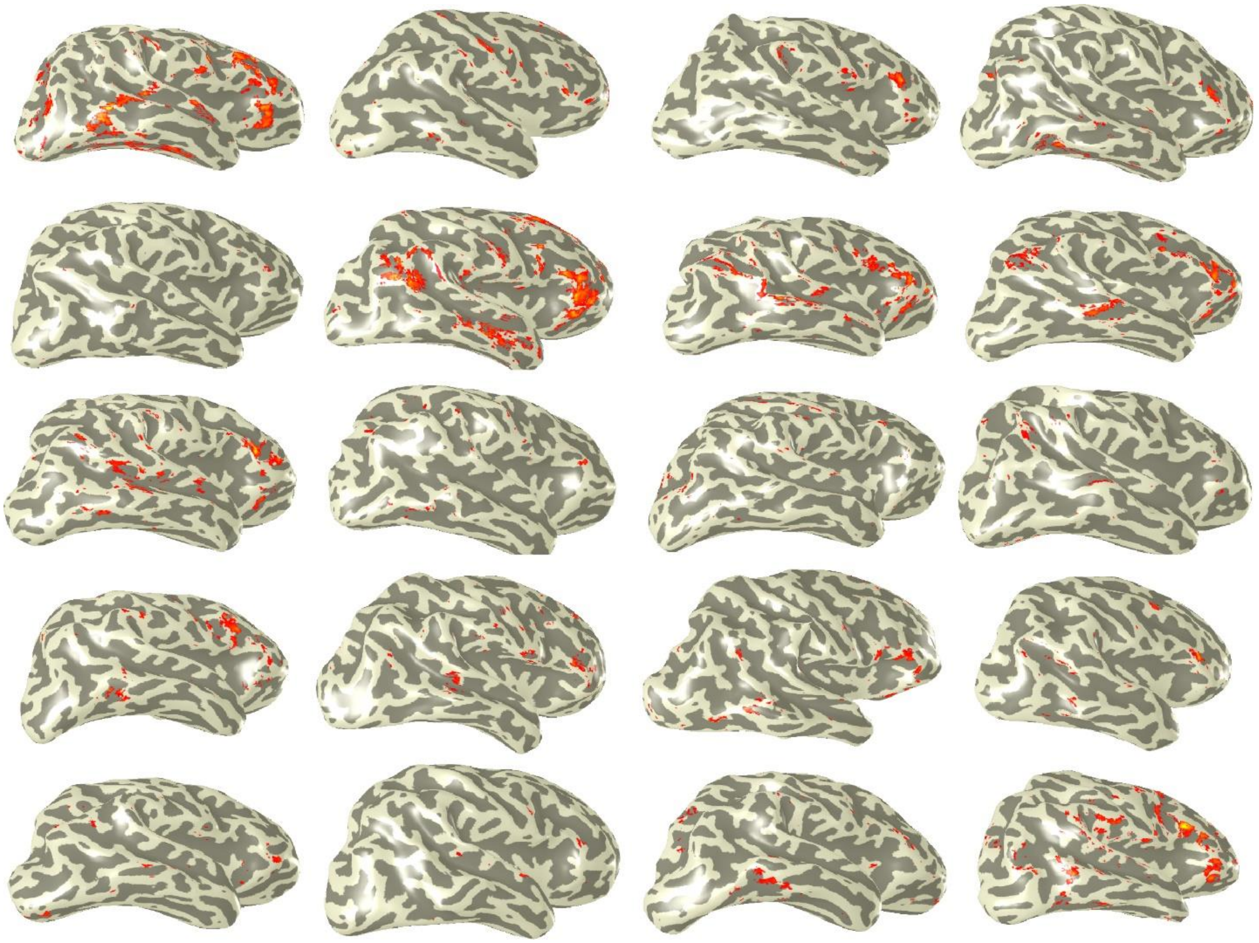

**Reading > Math+ Color (T≥3)**

**Supplementary Fig. 4. Multiple regions in the brain show higher responses to the reading task than the math and color tasks.** BOLD responses contrasting the reading task with the math and the color tasks (T≥3) are presented in the right hemisphere of all subjects. Subjects are sorted by their response accuracy in the math task, from highest (*top left*) to lowest (*bottom right*) mean performance.

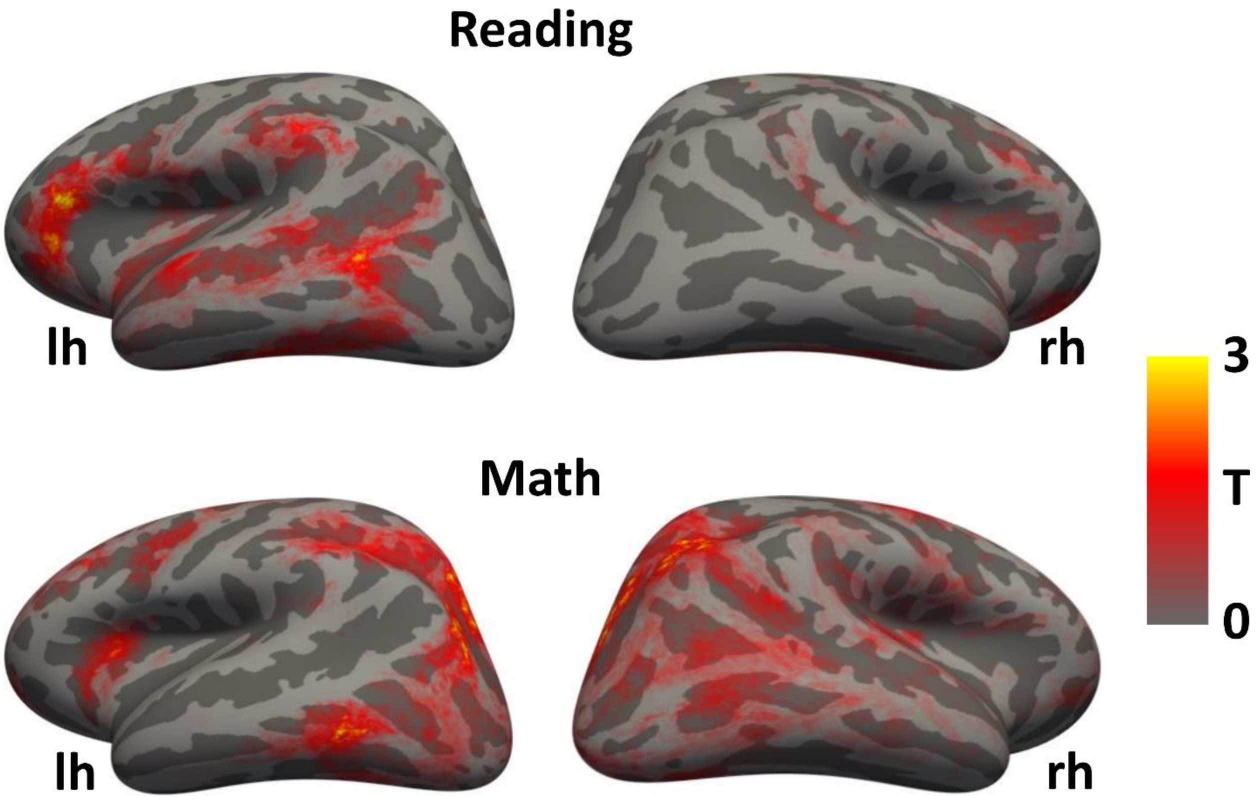


**Supplementary Fig. 5. On average, multiple regions in the brain show higher responses to the math or the reading task than the other tasks.** Parameter maps contrasting the reading task with the math and the color tasks (top) or the math task with the reading and the color task (bottom) were averaged across participants and are presented in the inflated left and right cortical surface of the FreeSurfer average brain.

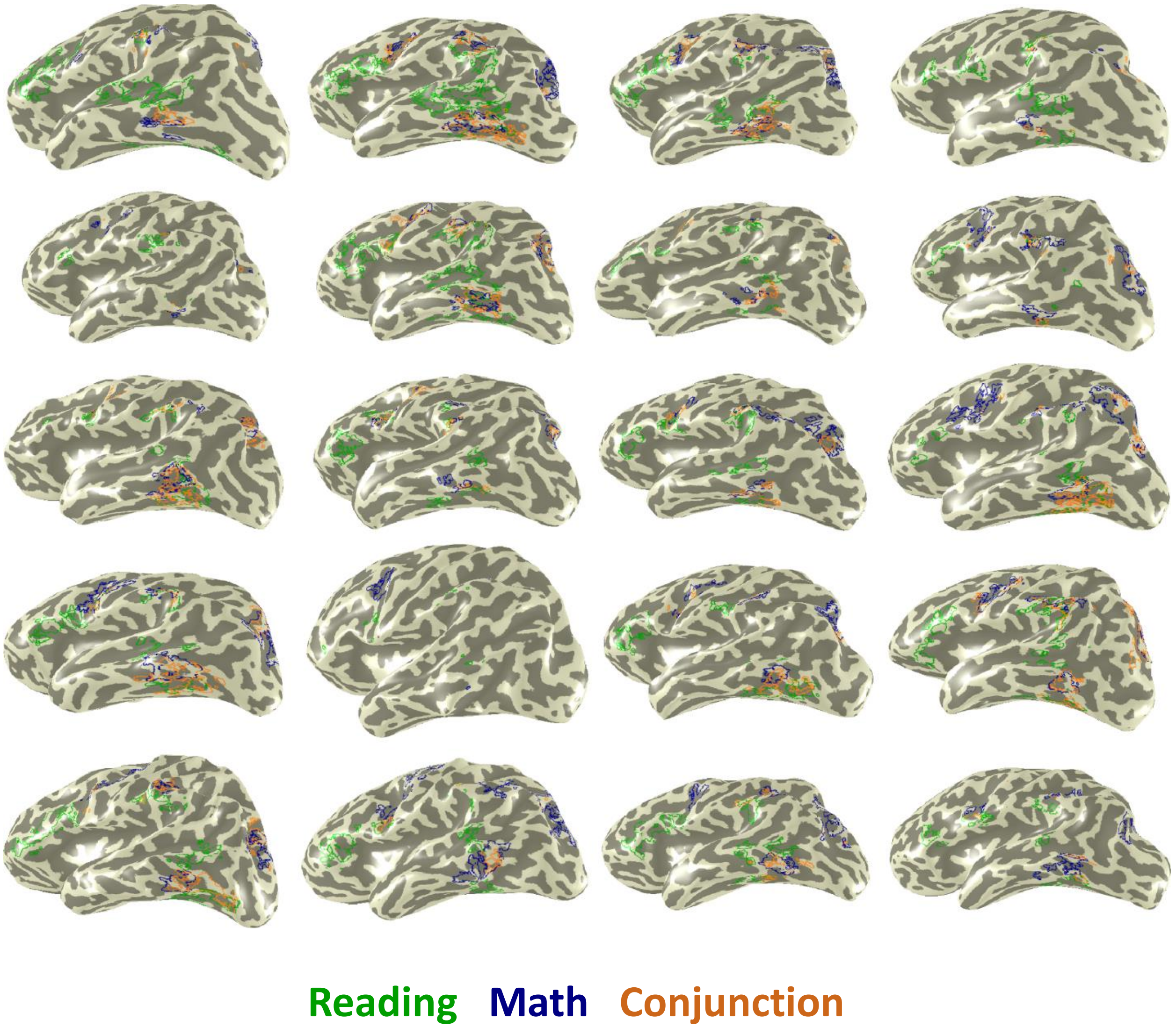


**Supplementary Fig. 6. Gray matter regions that respond more strongly to both math and reading than color are heavily overlapping with regions involved in math.** Regions were defined based on higher responses (T≥3) to reading compared to the other tasks (green outline), math compared to the other tasks (blue outline), or both math and reading compared to the color task (conjunction analysis, cyan outline) and are presented in the inflated left hemisphere of all participants.

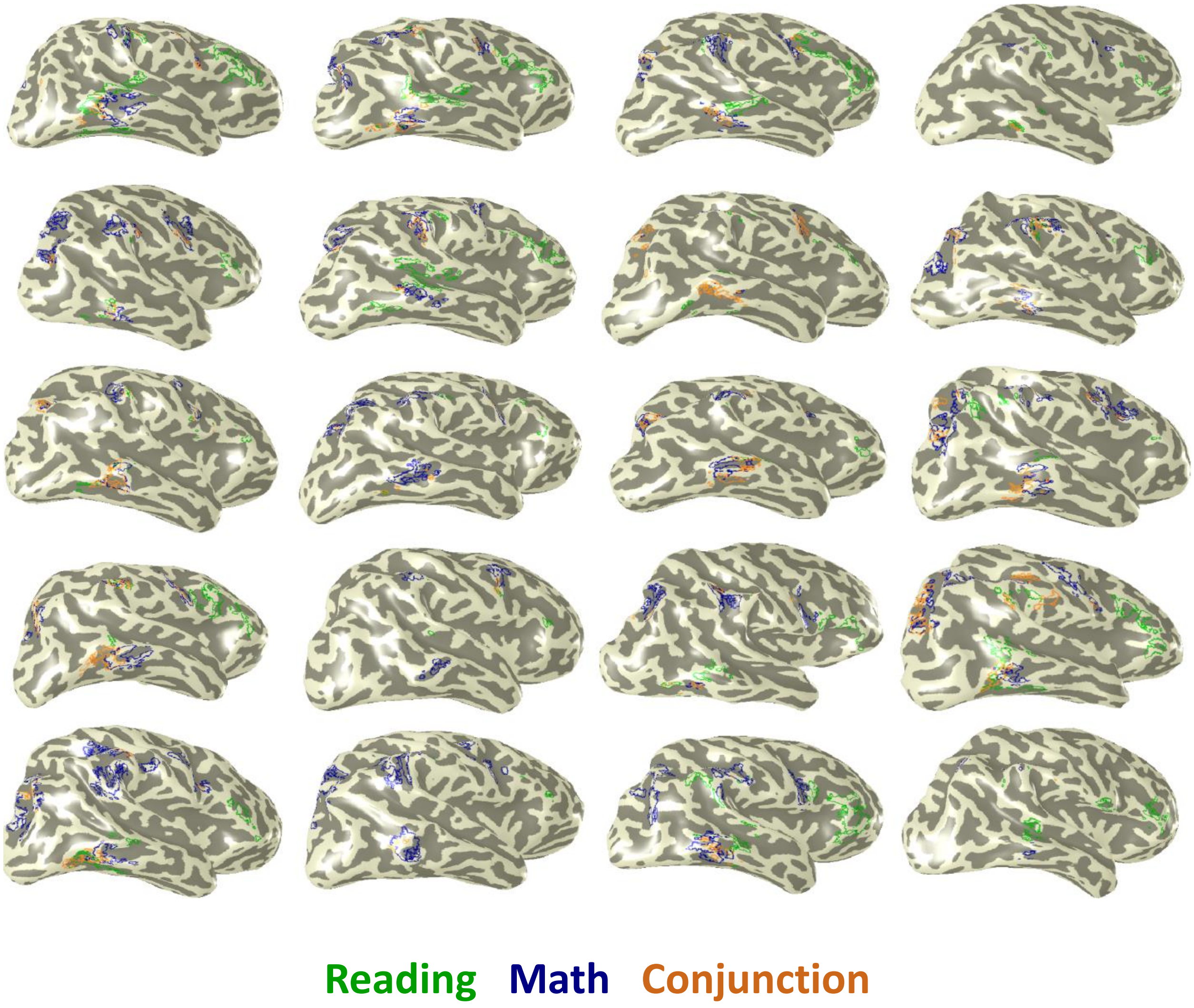


**Supplementary Fig. 7. Gray matter regions that respond more strongly to both math and reading than color are heavily overlapping with regions involved in math.** Regions were defined based on higher responses (T≥3) to reading compared to the other tasks (green outline), math compared to the other tasks (blue outline), or both math and reading compared to the color task (conjunction analysis, cyan outline) and are presented in the inflated right hemisphere of all participants.

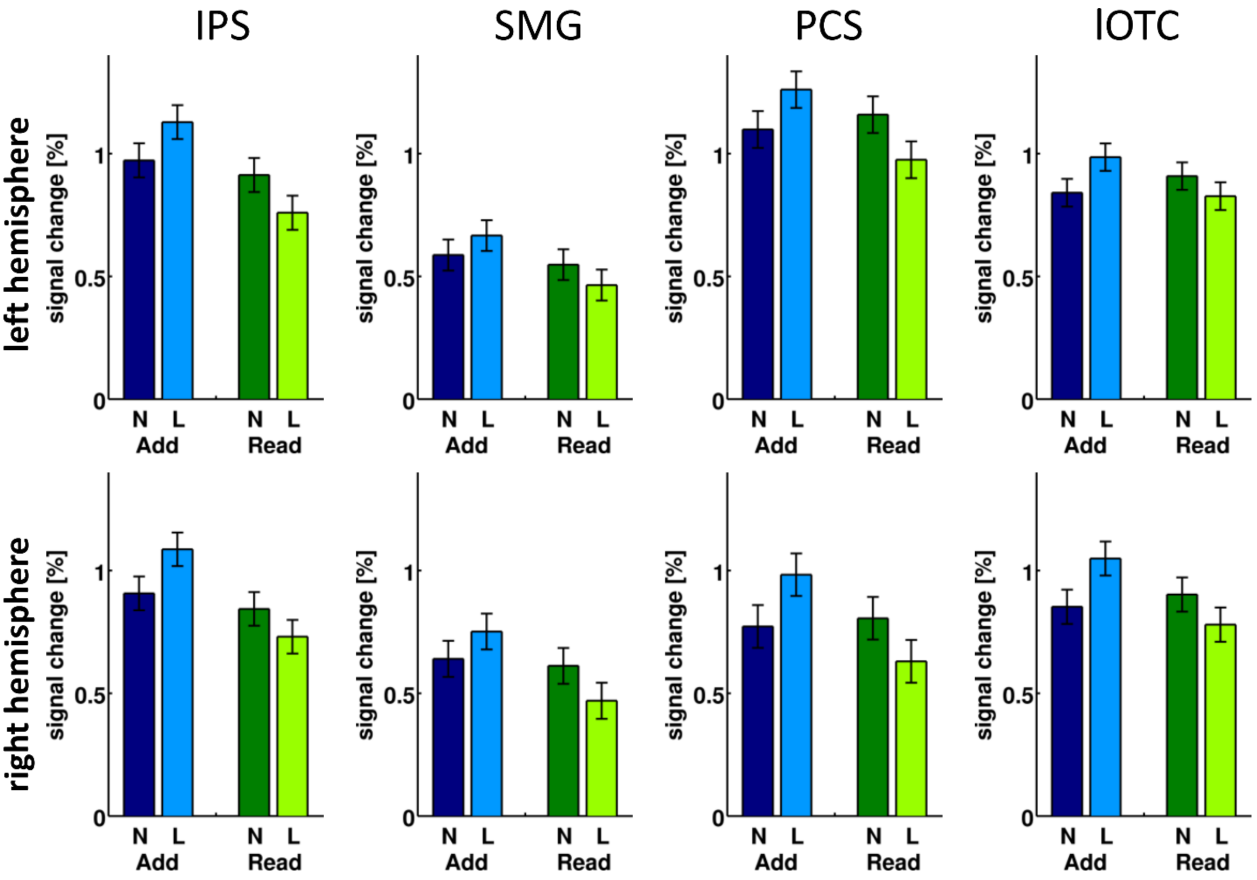


**Supplementary Fig. 8. Regions that show higher responses to math and reading than color prefer the math task over the reading task.** Response profile of regions identified in the conjunction analysis. All regions showed an interaction between task and stimulus (IPS: $F(1,14)=40.10$, $p<0.0001$, $\eta p^2=0.74$; PCS: $F(1,16)=49.07$, $p<0.0001$, $\eta p^2=0.75$; SMG: $F(1,16)=30.69$, $p<0.0001$, $\eta p^2=0.66$; lOTC: $F(1,17)=61.14$, $p<0.0001$, $\eta p^2=0.78$). The IPS, PCS and SMG, but not the lOTC regions, further showed a main effect of task, with higher responses in the adding task compared to the reding task (IPS: $F(1,14)=17.30$, $p=0.001$, $\eta p^2=0.55$; PCS: $F(1,16)=12.97$, $p-0.002$, $\eta p^2=0.45$; SMG: $F(1,16)=19.37$, $p=0.0004$, $\eta p^2=0.55$; lOTC: $F(1,17)=3.83$, $p=0.07$, $\eta p^2=0.18$). The PCS further showed a main effect of hemisphere, with higher responses in the left than the right hemisphere (PCS: $F(1,16)=13.43$, $p-0.002$, $\eta p^2=0.46$), while the SMG showed a weak interaction between stimulus and hemisphere $F(1,16)=4.70$, $p=0.05$, $\eta p^2=0.23$). No other main effects or interactions where observed. *Abbreviations*: IPS=intraparietal sulcus, SMG=supramarginal gyrus, PCS=precentral sulcus, lOTC=lateral occipito-temporal cortex.

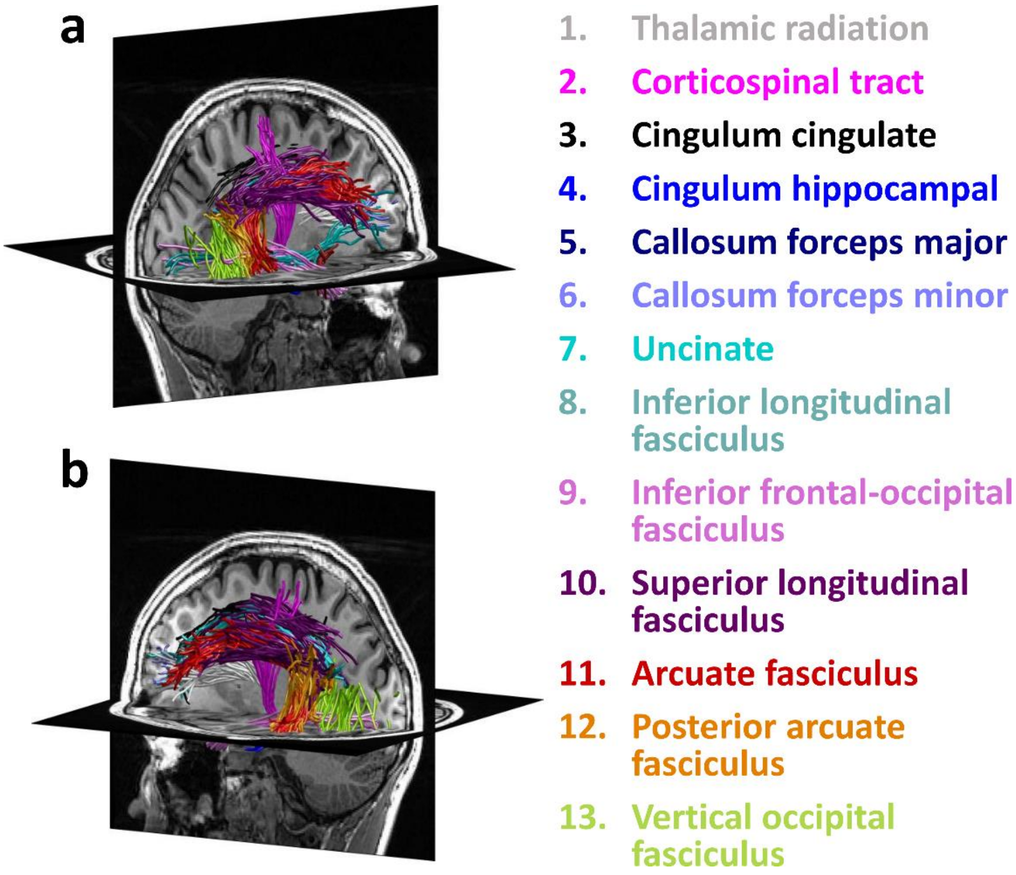


**Supplementary Fig. 9. Fascicles identified with the automatic fiber quantification toolbox (AFQ).** Fascicles were identified in both the left (a) and the right (b) hemispheres.

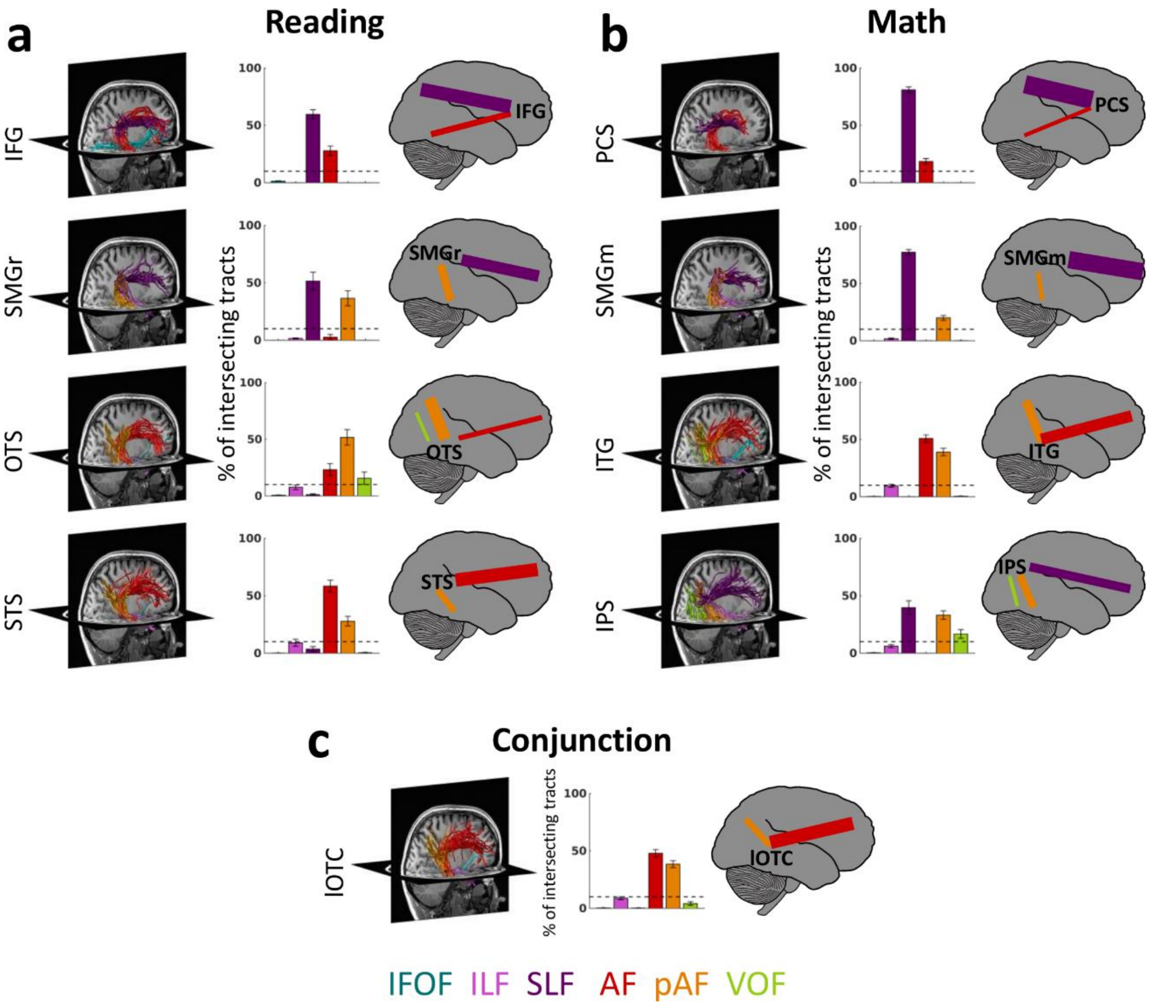


**Supplementary Fig. 10. Same as Fig. 2, but for the right hemisphere. Functionally-defined white matter tracts (fWMT) of reading and math related regions. (a)** Six fascicles (AF, SLF, pAF, VOF, ILF, and IFOF) contain ~90% of all fWMT of the fROIs identified in the reading task. **(b)** The same six fascicles also contain ~90% of all fWMT of the fROIs identified in the math task. **(c)** The conjunction fROI in the lOTC shows substantial connectivity with the AF and pAF. In **(a, b, c)**: *Left:* fWMT for each fROI in a representative subject's left hemisphere. The same subject is displayed in all panels; Fascicles are color coded in accordance with the legend at the bottom. *Middle*: Bar graphs showing what percentage of the fWMT is associated with each of the six fascicles. The graph shows the mean across subjects ± SEM. *Dashed horizontal line*: Line is placed at 10%, which was the cut-off used for the schematics in the right columns. *Right:* Schematic illustration of the fascicles associated with each fROI. The thickness of the lines is derived from the bar graph, showing the relative weight of each fascicle. *Abbreviations*: IFG=inferior frontal gyrus, PCS=precentral sulcus, SMGr=reading fROI in supramarginal gyrus, SMGm=math fROI in supramarginal gyrus, STS=superior temporal sulcus, ITG=inferior temporal gyrus, OTS=occipito-temporal sulcus, IPS=intraparietal sulcus, lOTC=lateral occipito-temporal cortex, IFOF=inferior fronto-occipital fasciculus, ILF=inferior longitudinal fasciculus, SLF=superior longitudinal fasciculus, AF=arcuate fasciculus, pAF=posterior arcuate fasciculus, VOF=vertical occipital fasciculus.

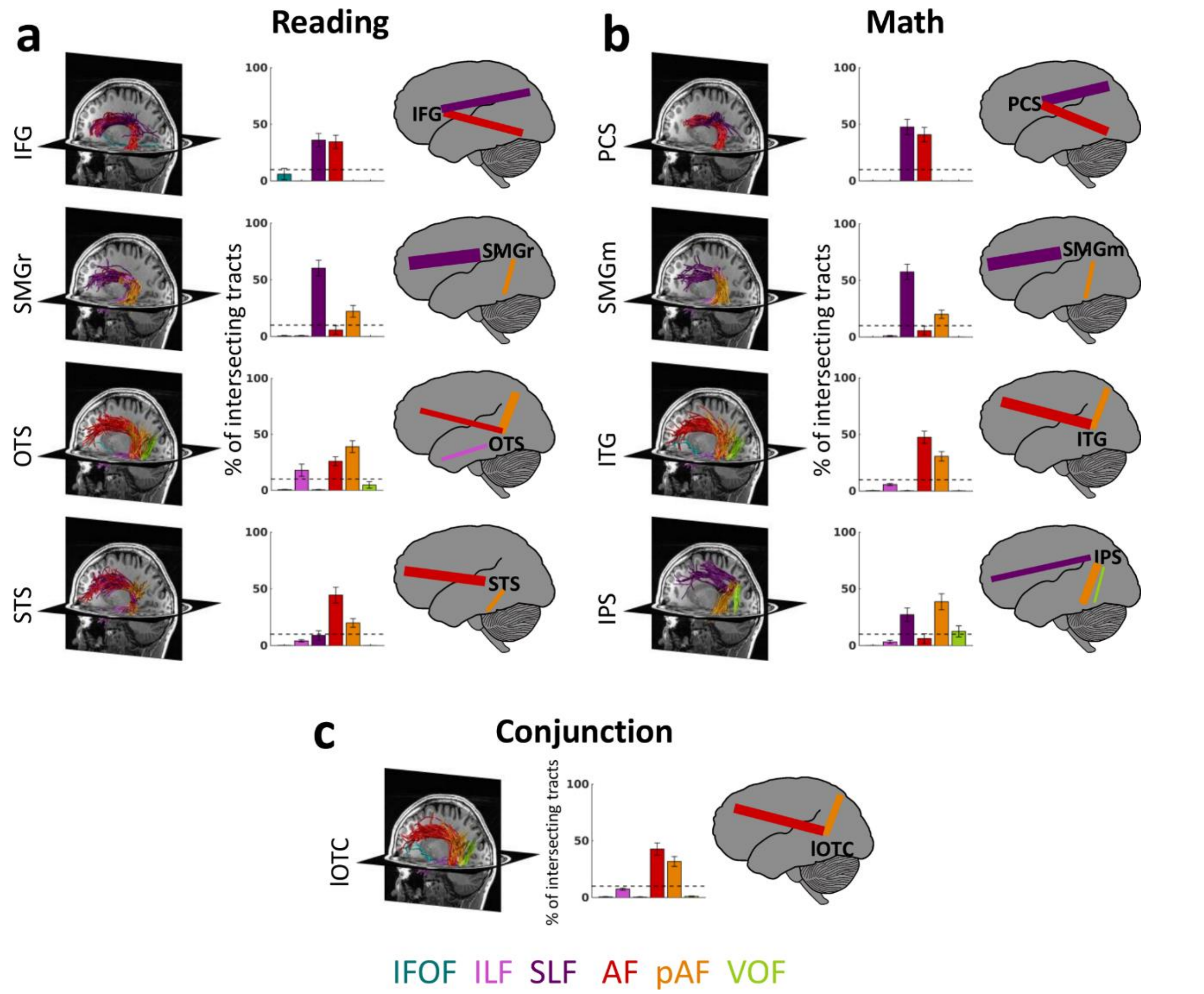


**Supplementary Fig. 11. Same as Fig. 2, but for functionally-defined white matter tracts (fWMT) identified with constant-size spherical ROIs of 7mm radius. (a)** Six fascicles (AF, SLF, pAF, VOF, ILF, and IFOF) contain ~90% of all fWMT of the fROIs identified in the reading task. **(b)** The same six fascicles also contain ~90% of all fWMT of the fROIs identified in the math task. **(c)** The conjunction fROI in the lOTC shows substantial connectivity with the AF and pAF. In **(a, b, c)**: *Left:* fWMT for each fROI in a representative subject's left hemisphere. The same subject is displayed in all panels; Fascicles are color coded in accordance with the legend at the bottom. *Middle*: Bar graphs showing what percentage of the fWMT is associated with each of the six fascicles. The graph shows the mean across subjects ± SEM. *Dashed horizontal line*: Line is placed at 10%, which was the cut-off used for the schematics in the right columns. *Right:* Schematic illustration of the fascicles associated with each fROI. The thickness of the lines is derived from the bar graph, showing the relative weight of each fascicle. *Abbreviations*: IFG=inferior frontal gyrus, PCS=precentral sulcus, SMGr=reading fROI in supramarginal gyrus, SMGm=math fROI in supramarginal gyrus, STS=superior temporal sulcus, ITG=inferior temporal gyrus, OTS=occipito-temporal sulcus, IPS=intraparietal sulcus, lOTC=lateral occipito-temporal cortex, IFOF=inferior fronto-occipital fasciculus, ILF=inferior longitudinal fasciculus, SLF=superior longitudinal fasciculus, AF=arcuate fasciculus, pAF=posterior arcuate fasciculus, VOF=vertical occipital fasciculus.

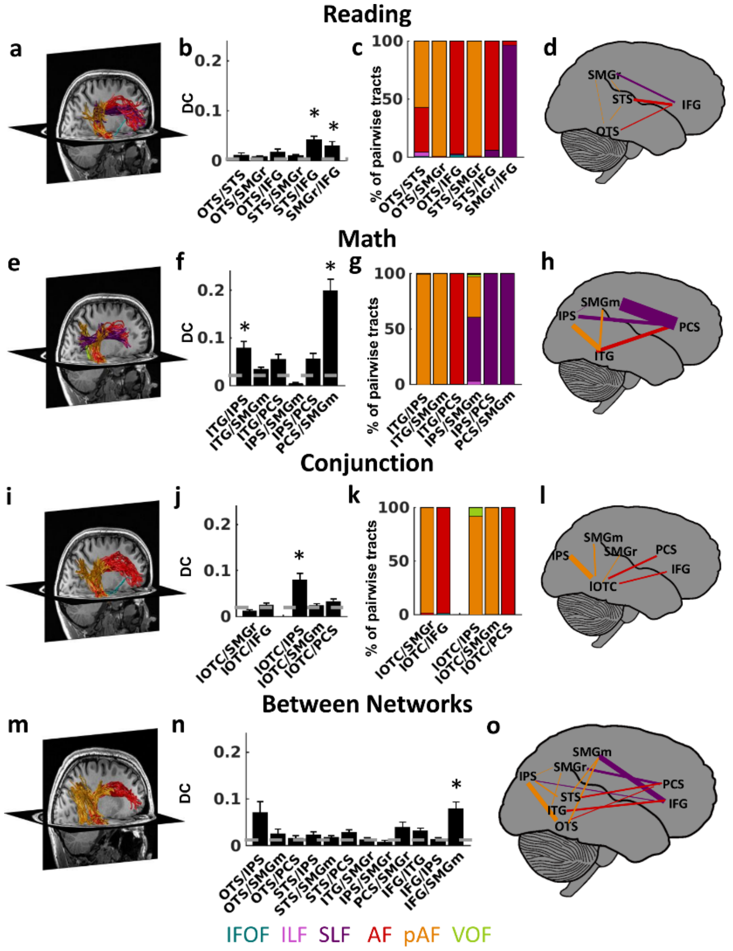


**Supplementary Fig. 12. Same as Fig. 3, but for the right hemisphere. Pairwise fWMT within and between the reading and math networks. (a-d)** Within-network connections of the reading network. **(e-h)** Within-network connections of the math network. **(i-j)** Connections of the conjunction fROI in the lOTC. **(m-o)** Between-network connections. *Left (a,e,i,m):* Pairwise white matter connections in a representative subject's left hemisphere. *Second from left (b,f,j,n):* Dice coefficient (DC) of pairwise connections, mean across subjects ± SEM. The DC quantifies the overlap in the fWMT of both fROIs: a DC of 1 indicates that all tracts that intersect with the first fROI also intersect with the second fROI, while a DC of 0 indicates no shared tracts. X-labels indicate the fROI pairing. *Dashed line:* Chance level DC estimated from the average connections to out of network fROIs in ventral temporal cortex that were activated maximally during the color task. *: DC is significantly higher than chance (significance level was Bonferroni adjusted). *Second from right in row 1-3 (c,g,k)*: The relative contribution of six fascicles to the pairwise connections (legend at bottom). X-labels indicate the fROI pairing. *Right (d,h,l,o):* Schematic illustration of the pairwise connections. *Line thickness* is scaled proportionally to the DC; *Color* indicates the fascicle with the highest relative contribution to pairwise connections. *Abbreviations:* IFG=inferior frontal gyrus, PCS=precentral sulcus, SMGr=reading fROI in supramarginal gyrus, SMGm=math fROI in supramarginal gyrus, STS=superior temporal sulcus, ITG=inferior temporal gyrus, OTS=occipito-temporal sulcus, IPS=intraparietal sulcus, lOTC=lateral occipito-temporal cortex, IFOF=inferior fronto-occipital fasciculus, ILF=inferior longitudinal fasciculus, SLF=superior longitudinal fasciculus, AF=arcuate fasciculus, pAF=posterior arcuate fasciculus, VOF=vertical occipital fasciculus.

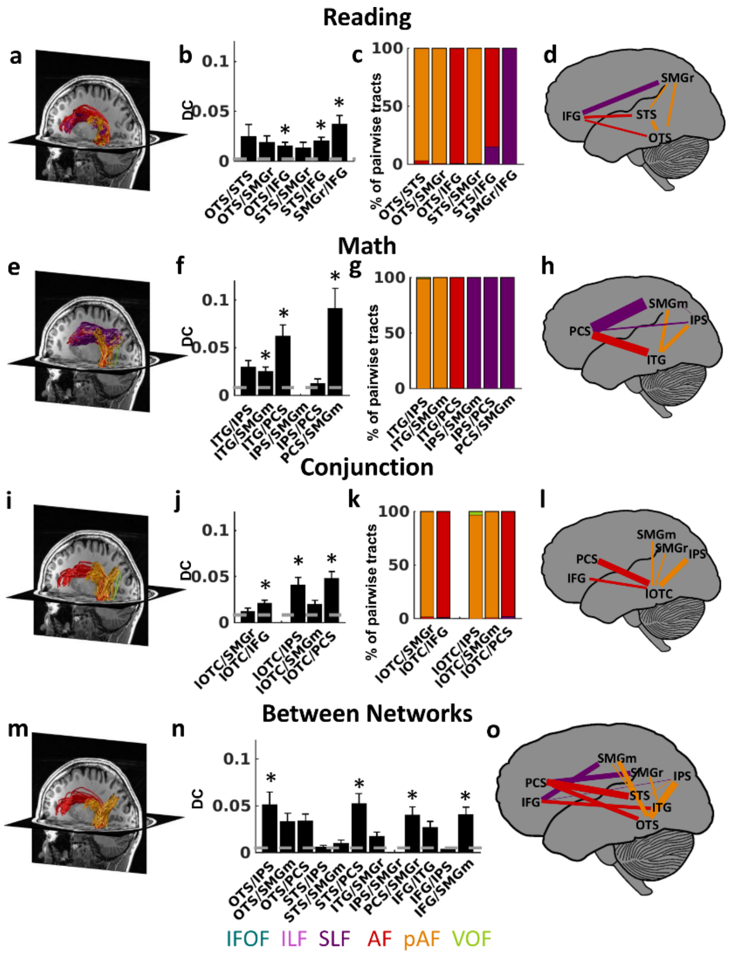


**Supplementary Fig. 13. Same as Fig. 3, but for fWMTs identified with constant-size spherical ROIs of 7mm radius. (a-d)** Within-network connections of the reading network. **(e-h)** Within-network connections of the math network. **(i-j)** Connections of the conjunction fROI in the lOTC. **(m-o)** Between-network connections. *Left (a,e,i,m):* Pairwise white matter connections in a representative subject's left hemisphere. *Second from left (b,f,j,n):* Dice coefficient (DC) of pairwise connections, mean across subjects ± SEM. The DC quantifies the overlap in the fWMT of both fROIs: a DC of 1 indicates that all tracts that intersect with the first fROI also intersect with the second fROI, while a DC of 0 indicates no shared tracts. X-labels indicate the fROI pairing. *Dashed line:* Chance level DC estimated from the average connections to out of network fROIs in ventral temporal cortex that were activated maximally during the color task. *: DC is significantly higher than chance (significance level was Bonferroni adjusted). *Second from right in row 1-3 (c,g,k)*: The relative contribution of six fascicles to the pairwise connections (legend at bottom). X-labels indicate the fROI pairing. *Right (d,h,l,o):* Schematic illustration of the pairwise connections. *Line thickness* is scaled proportionally to the DC; *Color* indicates the fascicle with the highest relative contribution to pairwise connections. *Abbreviations:* IFG=inferior frontal gyrus, PCS=precentral sulcus, SMGr=reading fROI in supramarginal gyrus, SMGm=math fROI in supramarginal gyrus, STS=superior temporal sulcus, ITG=inferior temporal gyrus, OTS=occipito-temporal sulcus, IPS=intraparietal sulcus, lOTC=lateral occipito-temporal cortex, IFOF=inferior fronto-occipital fasciculus, ILF=inferior longitudinal fasciculus, SLF=superior longitudinal fasciculus, AF=arcuate fasciculus, pAF=posterior arcuate fasciculus, VOF=vertical occipital fasciculus.

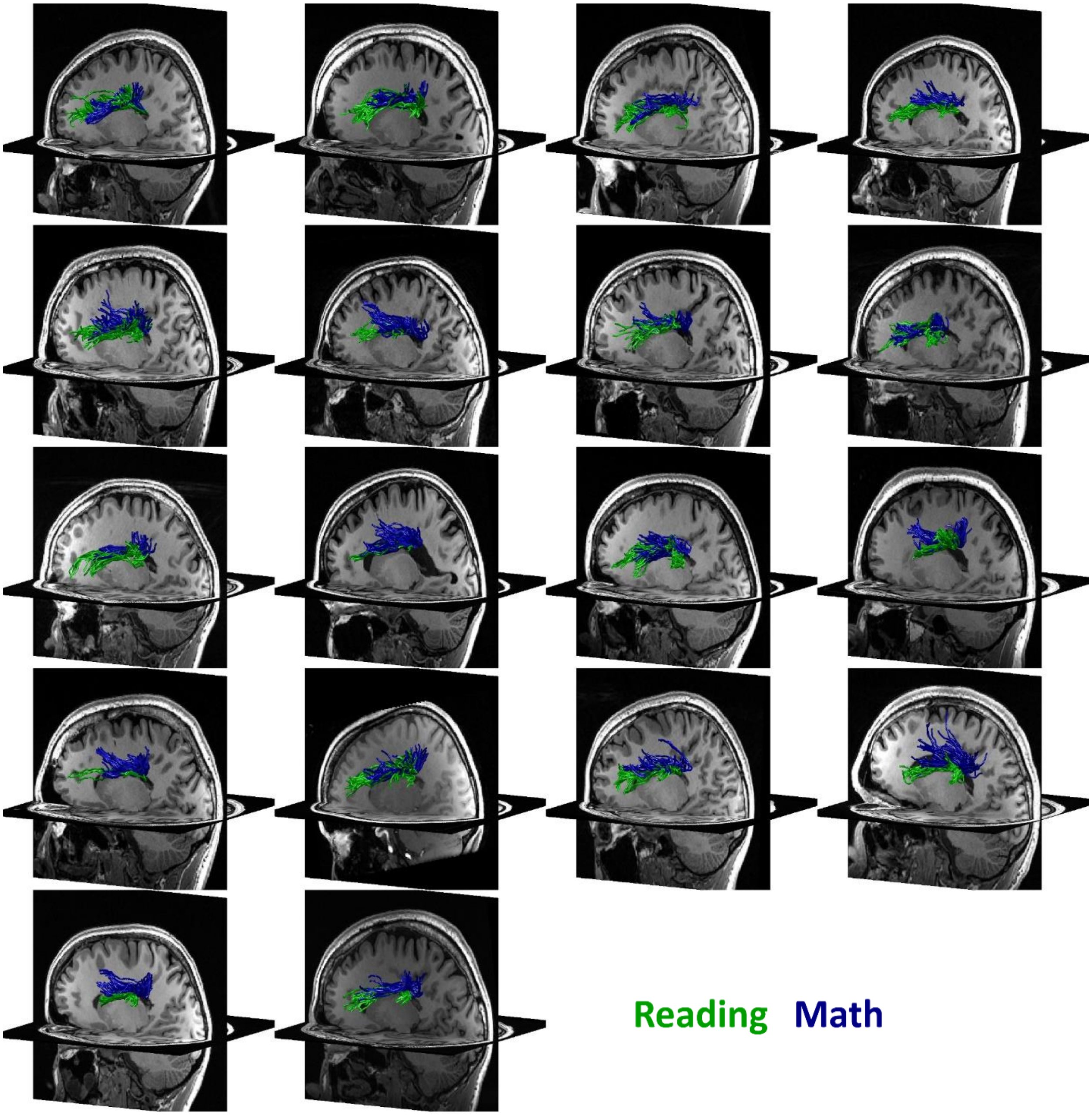


**Supplementary Fig. 14. Pairwise connections within the reading and the math networks are segregated and parallel in the SLF.** SLF tracts connecting IFG and SMG in the reading network (green) and PCS and SMG in the math network (blue) are presented in the left hemisphere of all individual subjects. Subjects show spatial segregation of these tracts. *Abbreviations*: IFG=inferior frontal gyrus, PCS=precentral sulcus, SMG=supramarginal gyrus, SLF=superior longitudinal fasciculus.

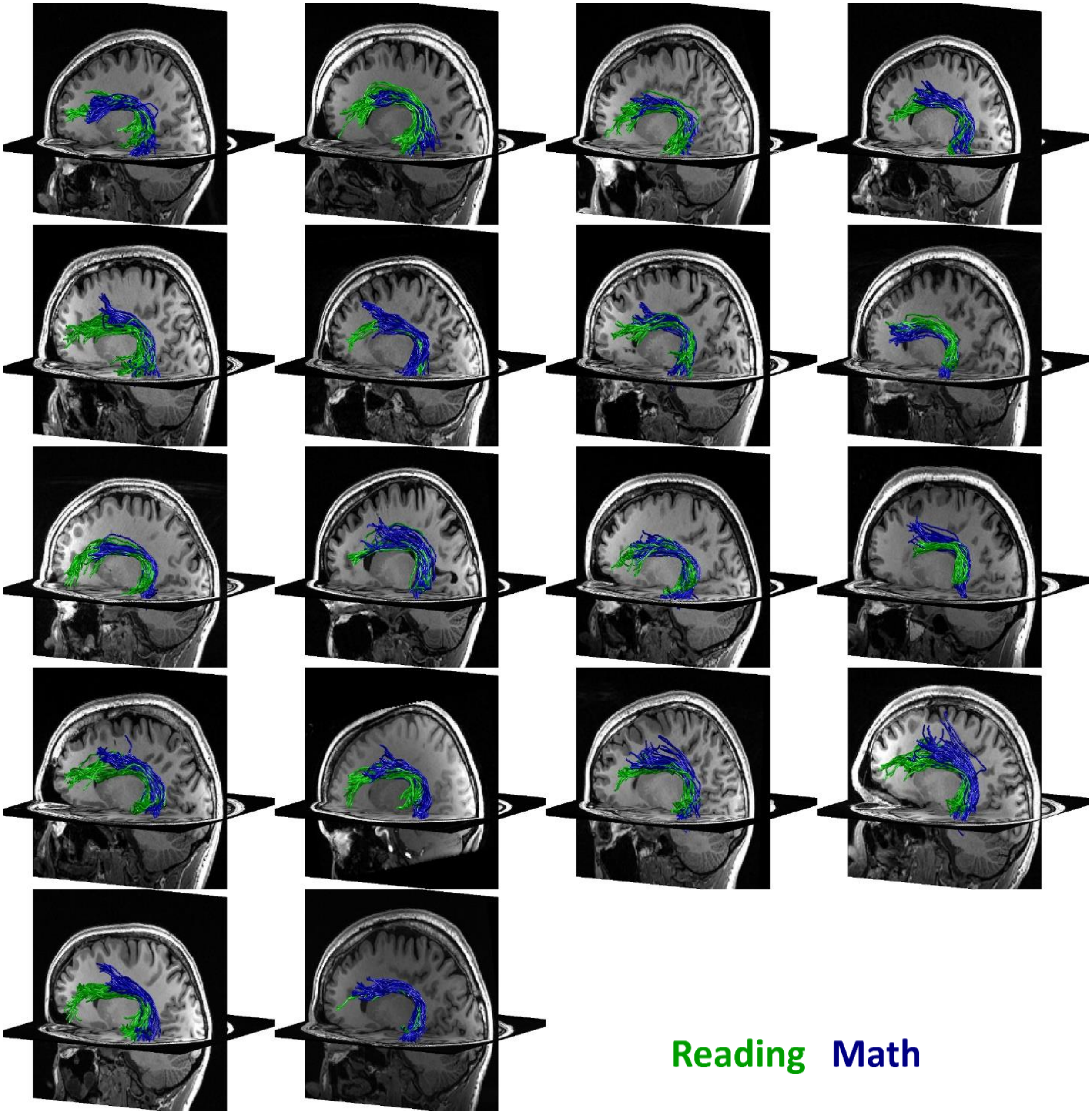


**Supplementary Fig. 15. Pairwise connections within the reading and the math networks are segregated and parallel in the AF.** AF tracts connecting IFG and STS in the reading network (green) and PCS and ITG in the math network (blue) are presented in the left hemisphere of all individual subjects. Subjects show spatial segregation of these tracts. *Abbreviations:* IFG=inferior frontal gyrus, PCS=precentral sulcus, STS=superior temporal sulcus, ITG=inferior temporal gyrus, SLF=superior longitudinal fasciculus.

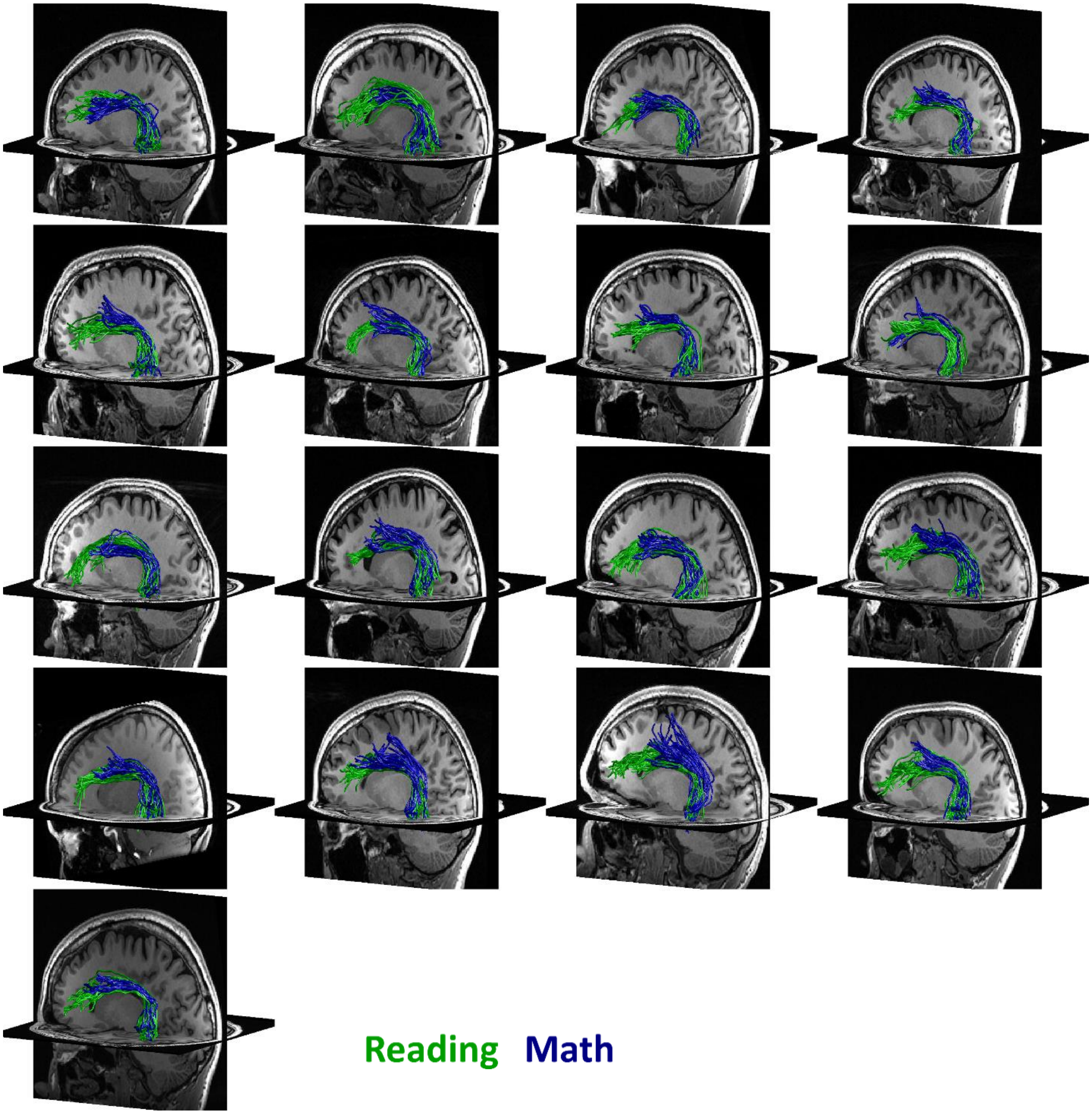


**Supplementary Fig. 16. Segregated tracts between the lOTC and frontal fROIs associated with math and reading, respectively.** AF tracts connecting the lOTC conjunction fROI with the IFG in the reading network (green) and the PCS in the math network (blue) are shown in the left hemisphere of all individual subjects. Subjects show spatial segregation of these tracts. *Abbreviations:* IFG=inferior frontal gyrus, PCS=precentral sulcus, lOTC=lateral occipito-temporal cortex.

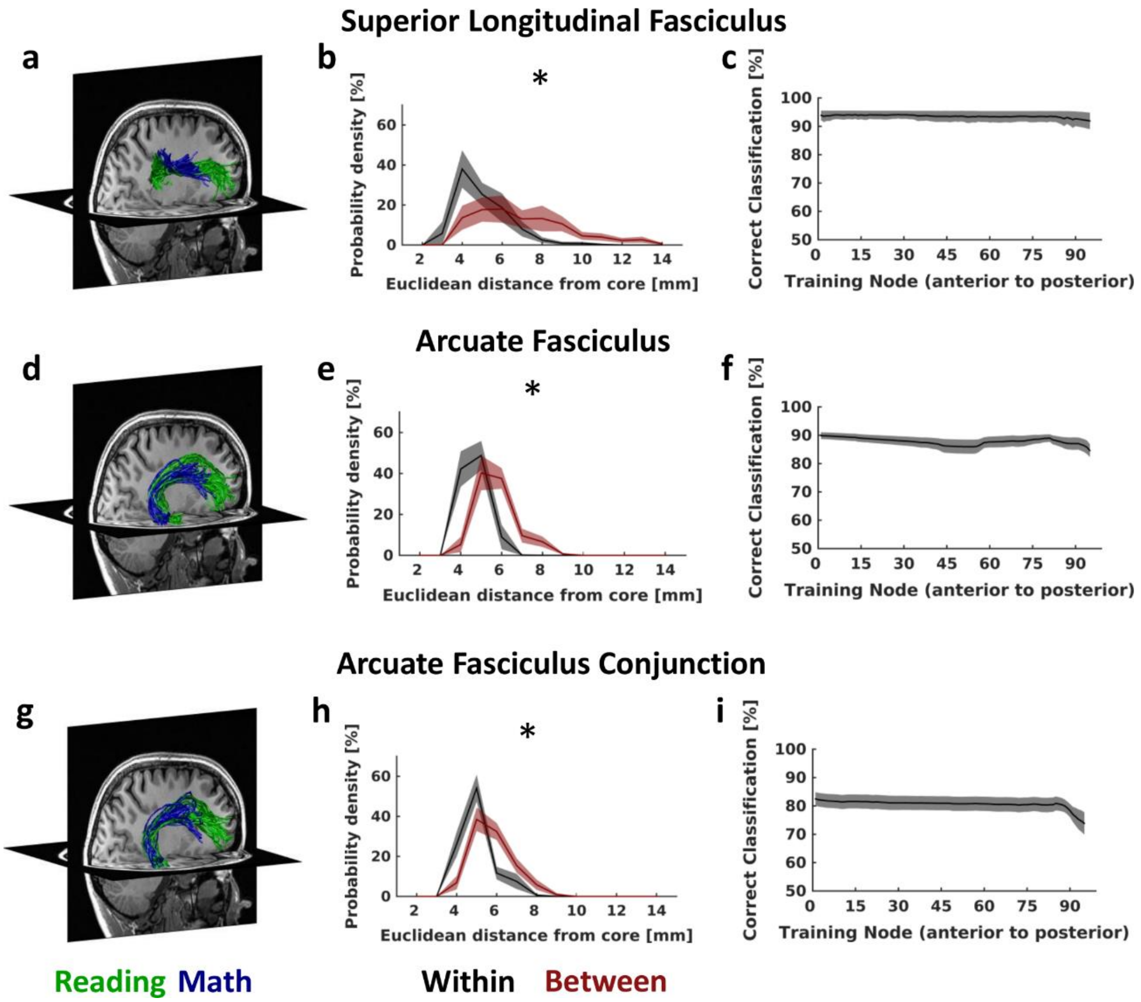


**Supplementary Fig. 17. Same as Fig. 4, but for the right hemisphere. Pairwise connections of the reading and the math networks are segregated and parallel in the SLF and the AF. (a-c)**: SLF tracts connecting IFG and SMGr in the reading network and PCS and SMGm in the math network. **(d-f)**: AF tracts connecting the IFG and STS in the reading network and the PCS and ITG in the math network. **(g-h)**: AF tracts connecting the lOTC fROI identified in the conjunction analysis with both the IFG in the reading network and the PCS in the math network. **(a,d,g)**: Math (blue) and reading (green) tracts of the SLF and AF in a representative individual subject showing the spatial segregation of these tracts. **(b,e,h)**: Euclidean distance in mm (derived from x,y,z coordinates) of all tracts relative to the core (mean) tract, within-network (black) and between-network (maroon). The distance was calculated across all tracts; the plot shows the mean across all nodes ±SEM. * Distributions differ significantly, $p<0.05$. **(c,f,i)**: Performance of a linear SVM classifying math and reading tracts within the SLF and AF based on their spatial location. Data show mean classification accuracy across nodes ±SEM. *Abbreviations:* IFG=inferior frontal gyrus, PCS=precentral sulcus, SMGr=reading fROI in supramarginal gyrus, SMGm=math fROI in supramarginal gyrus, STS=superior temporal sulcus, ITG=inferior temporal gyrus, lOTC=lateral occipito-temporal cortex, AF=arcuate fasciculus, SLF=superior longitudinal fasciculus.

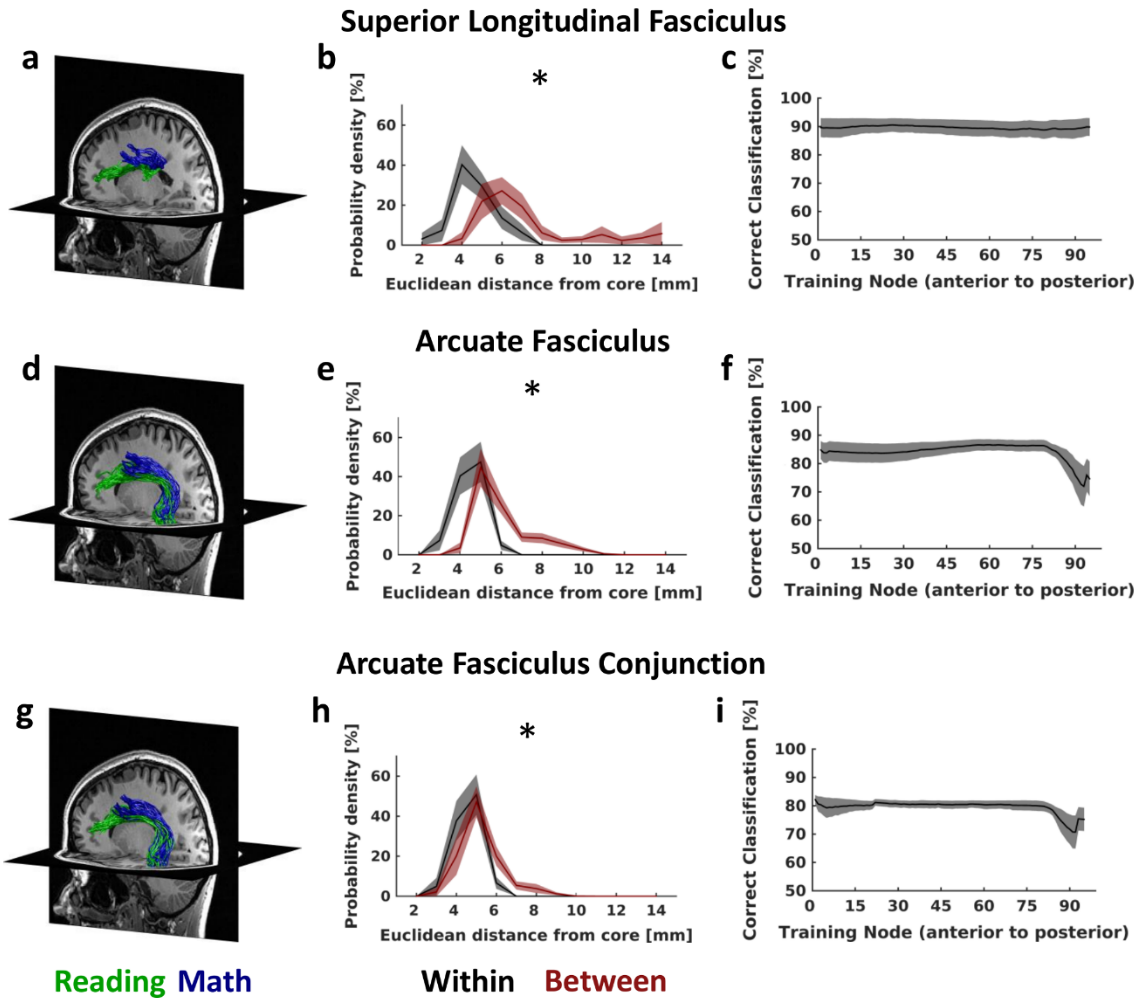


**Supplementary Fig. 18. Same as Fig. 4, but for tracts identified with constant-size spherical ROIs of 7mm radius. (a-c)**: SLF tracts connecting IFG and SMGr in the reading network and PCS and SMGm in the math network. **(d-f)**: AF tracts connecting the IFG and STS in the reading network and the PCS and ITG in the math network. **(g-h)**: AF tracts connecting the lOTC fROI identified in the conjunction analysis with both the IFG in the reading network and the PCS in the math network. **(a,d,g)**: Math (blue) and reading (green) tracts of the SLF and AF in a representative individual subject showing the spatial segregation of these tracts. **(b,e,h)**: Euclidean distance in mm (derived from x,y,z coordinates) of all tracts relative to the core (mean) tract, within-network (black) and between-network (maroon). The distance was calculated across all tracts; the plot shows the mean across all nodes ±SEM. * Distributions differ significantly, $p<0.05$. **(c,f,i)**: Performance of a linear SVM classifying math and reading tracts within the SLF and AF based on their spatial location. Data show mean classification accuracy across nodes ±SEM. *Abbreviations:* IFG=inferior frontal gyrus, PCS=precentral sulcus, SMGr=reading fROI in supramarginal gyrus, SMGm=math fROI in supramarginal gyrus, STS=superior temporal sulcus, ITG=inferior temporal gyrus, lOTC=lateral occipito-temporal cortex, AF=arcuate fasciculus, SLF=superior longitudinal fasciculus.

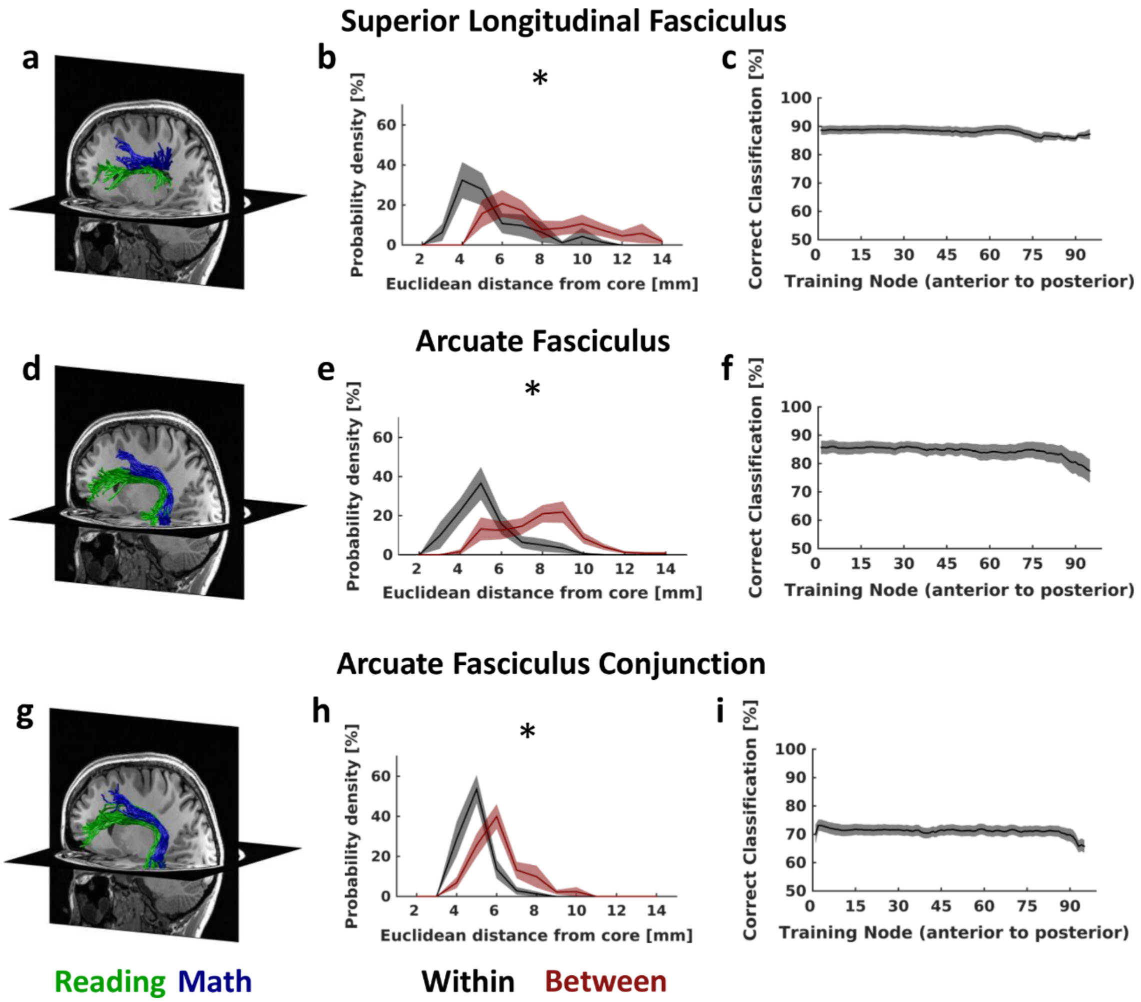


**Supplementary Fig. 19. Same as Fig. 4, but for fWMT identified using fROI to fROI tractography. (a-c)**: SLF tracts connecting IFG and SMGr in the reading network and PCS and SMGm in the math network. **(d-f)**: AF tracts connecting the IFG and STS in the reading network and the PCS and ITG in the math network. **(g-h)**: AF tracts connecting the lOTC fROI identified in the conjunction analysis with both the IFG in the reading network and the PCS in the math network. **(a,d,g)**: Math (blue) and reading (green) tracts of the SLF and AF in a representative individual subject showing the spatial segregation of these tracts. **(b,e,h)**: Euclidean distance in mm (derived from x,y,z coordinates) of all tracts relative to the core (mean) tract, within-network (black) and between-network (maroon). The distance was calculated across all tracts; the plot shows the mean across all nodes ±SEM. * Distributions differ significantly, $p<0.05$. **(c,f,i)**: Performance of a linear SVM classifying math and reading tracts within the SLF and AF based on their spatial location. Data show mean classification accuracy across nodes ±SEM. *Abbreviations:* IFG=inferior frontal gyrus, PCS=precentral sulcus, SMGr=reading fROI in supramarginal gyrus, SMGm=math fROI in supramarginal gyrus, STS=superior temporal sulcus, ITG=inferior temporal gyrus, lOTC=lateral occipito-temporal cortex, AF=arcuate fasciculus, SLF=superior longitudinal fasciculus.

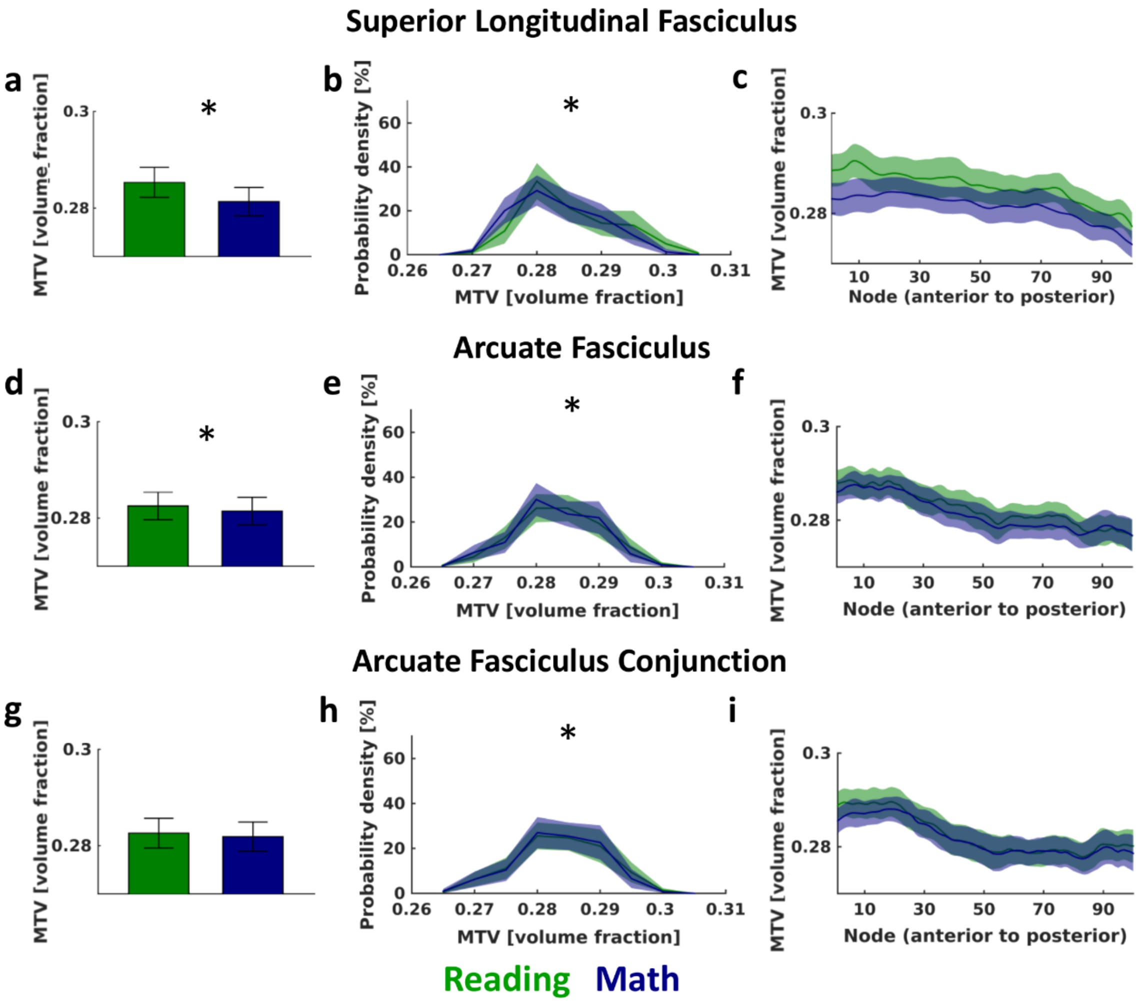


**Supplementary Fig. 20. Left hemisphere SLF and AF tracts associated with reading show higher macromolecular tissue volume fraction (MTV) than those associated with math. (a-c)**: MTV measurements for SLF tracts connecting IFG and SMGr in the reading network (green) and PCS and SMGm in the math network (blue). **(d-f)**: MTV measurements for AF tracts connecting the IFG and STS in the reading network (green) and the PCS and ITG in the math network (blue). **(g-i)**: MTV measurements for AF tracts connecting the lOTC conjunction fROI with the IFG in the reading network (green) and the PCS in the math network (blue). Left **(a,d,g)**: Average MTV for reading- and math-related tracts in the SLF and the AF. Bar graph shows mean across subjects ± SEM. *: MTV for math and reading related tracts differs significantly, $p<0.05$. Middle **(b,e,h)**: Distribution of MTV values across all tracts. Distributions were calculated within each subject and node; the plot shows the mean across all nodes ± SEM. *: Distributions differ significantly, $p<0.05$. Right **(c,f,i)**: Average MTV for reading- and math-related tracts along the SLF and the AF. Line graph shows mean across subjects ± SEM. *Abbreviations:* IFG=inferior frontal gyrus, PCS=precentral sulcus, SMGr=reading fROI in supramarginal gyrus, SMGm=math fROI in supramarginal gyrus, STS=superior temporal sulcus, ITG=inferior temporal gyrus, lOTC=lateral occipito-temporal cortex, SLF=superior longitudinal fasciculus, AF=arcuate fasciculus.

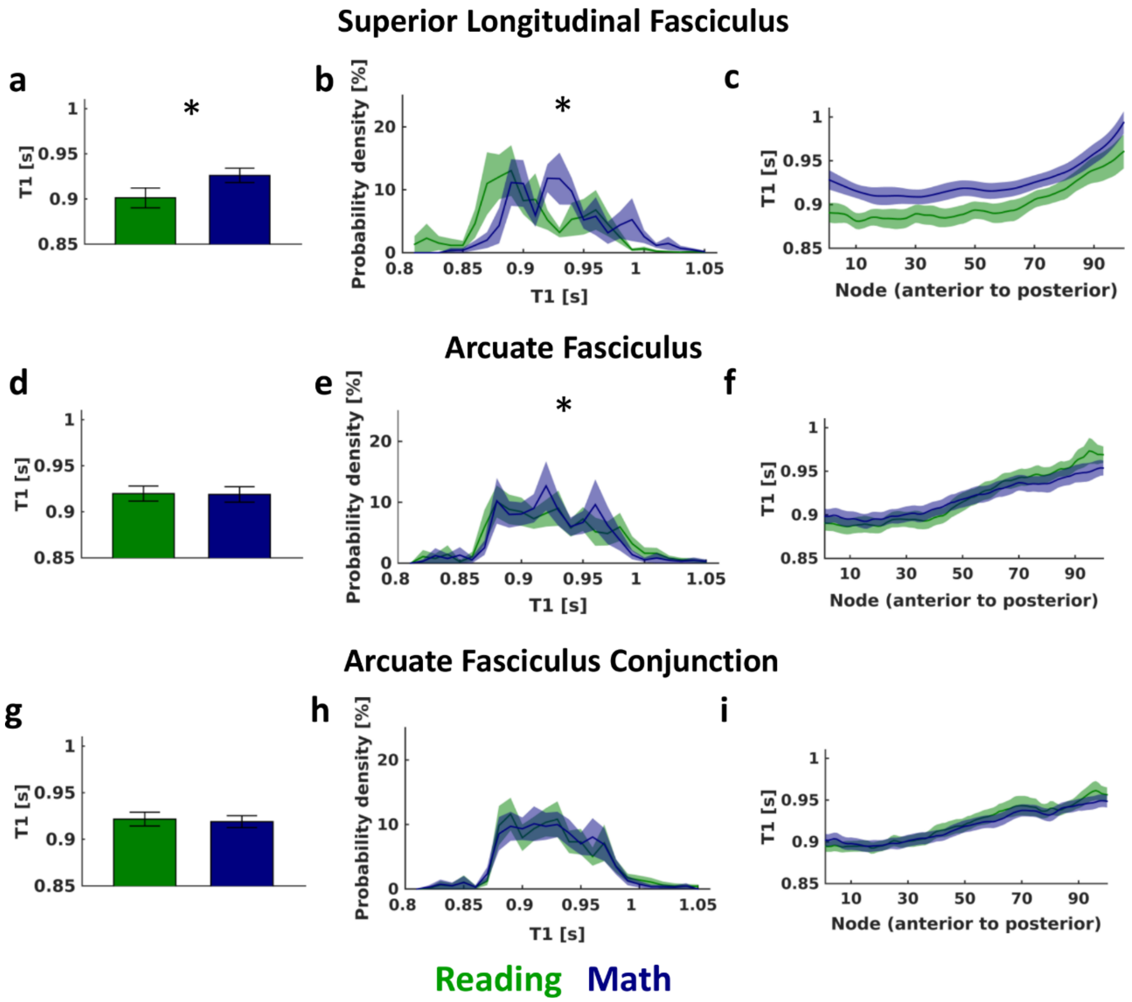


**Supplementary Fig. 21. Same as Fig. 5, but for the right hemisphere. Tracts associated with reading show faster proton relaxation time ($T_1$) than those associated with math. (a-c)**: $T_1$ measurements for SLF tracts connecting IFG and SMGr in the reading network (green) and PCS and SMGm in the math network (blue). **(d-f)**: $T_1$ measurements for AF tracts connecting the IFG and STS in the reading network (green) and the PCS and ITG in the math network (blue). **(g-i)**: $T_1$ measurements for AF tracts connecting the lOTC conjunction fROI with the IFG in the reading network (green) and the PCS in the math network (blue). Left **(a,d,g)**: Average $T_1$ for reading- and math-related tracts in the SLF and the AF. Bar graph shows mean across subjects ± SEM. *: $T_1$ for math and reading related tracts differs significantly, $p<0.05$. Middle **(b,e,h)**: Distribution of $T_1$ values across all tracts. Distributions were calculated within each subject and node; the plot shows the mean across all nodes ± SEM. *: Distributions differ significantly, $p<0.05$. Right **(c,f,i)**: Average $T_1$ for reading- and math-related tracts along the SLF and the AF. Line graph shows mean across subjects ± SEM. *Abbreviations:* IFG=inferior frontal gyrus, PCS=precentral sulcus, SMGr=reading fROI in supramarginal gyrus, SMGm=math fROI in supramarginal gyrus, STS=superior temporal sulcus, ITG=inferior temporal gyrus, lOTC=lateral occipito-temporal cortex, SLF=superior longitudinal fasciculus, AF=arcuate fasciculus.

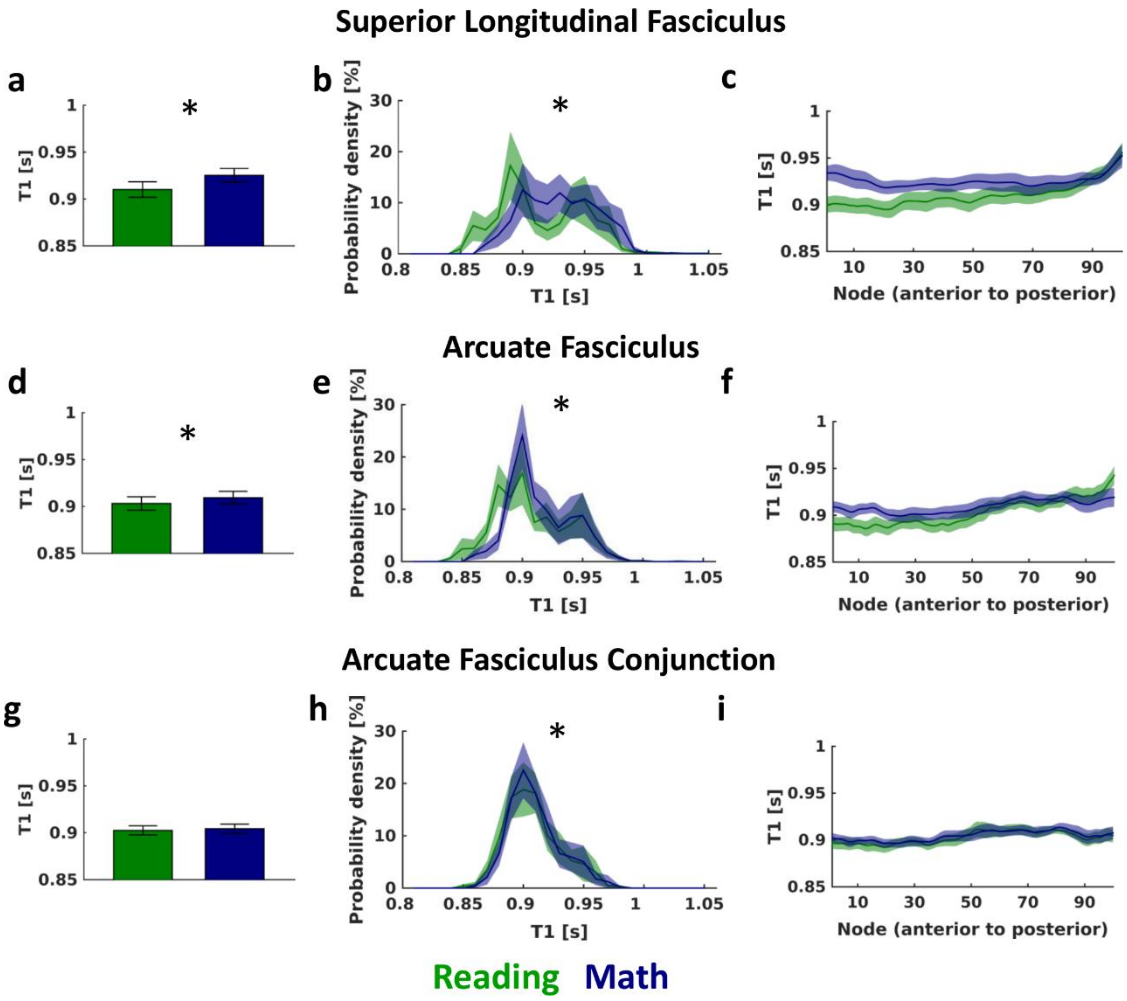


**Supplementary Fig. 22. Same as Fig. 5, but for tracts identified using constant-size spherical ROIs of 7mm radius. (a-c)**: $T_1$ measurements for SLF tracts connecting IFG and SMGr in the reading network (green) and PCS and SMGm in the math network (blue). **(d-f)**: $T_1$ measurements for AF tracts connecting the IFG and STS in the reading network (green) and the PCS and ITG in the math network (blue). **(g-i)**: $T_1$ measurements for AF tracts connecting the lOTC conjunction fROI with the IFG in the reading network (green) and the PCS in the math network (blue). Left **(a,d,g)**: Average $T_1$ for reading- and math-related tracts in the SLF and the AF. Bar graph shows mean across subjects ± SEM. *: $T_1$ for math and reading related tracts differs significantly, $p<0.05$. Middle **(b,e,h)**: Distribution of $T_1$ values across all tracts. Distributions were calculated within each subject and node; the plot shows the mean across all nodes ± SEM. *: Distributions differ significantly, $p<0.05$. Right **(c,f,i)**: Average $T_1$ for reading- and math-related tracts along the SLF and the AF. Line graph shows mean across subjects ± SEM. *Abbreviations:* IFG=inferior frontal gyrus, PCS=precentral sulcus, SMGr=reading fROI in supramarginal gyrus, SMGm=math fROI in supramarginal gyrus, STS=superior temporal sulcus, ITG=inferior temporal gyrus, lOTC=lateral occipito-temporal cortex, SLF=superior longitudinal fasciculus, AF=arcuate fasciculus.

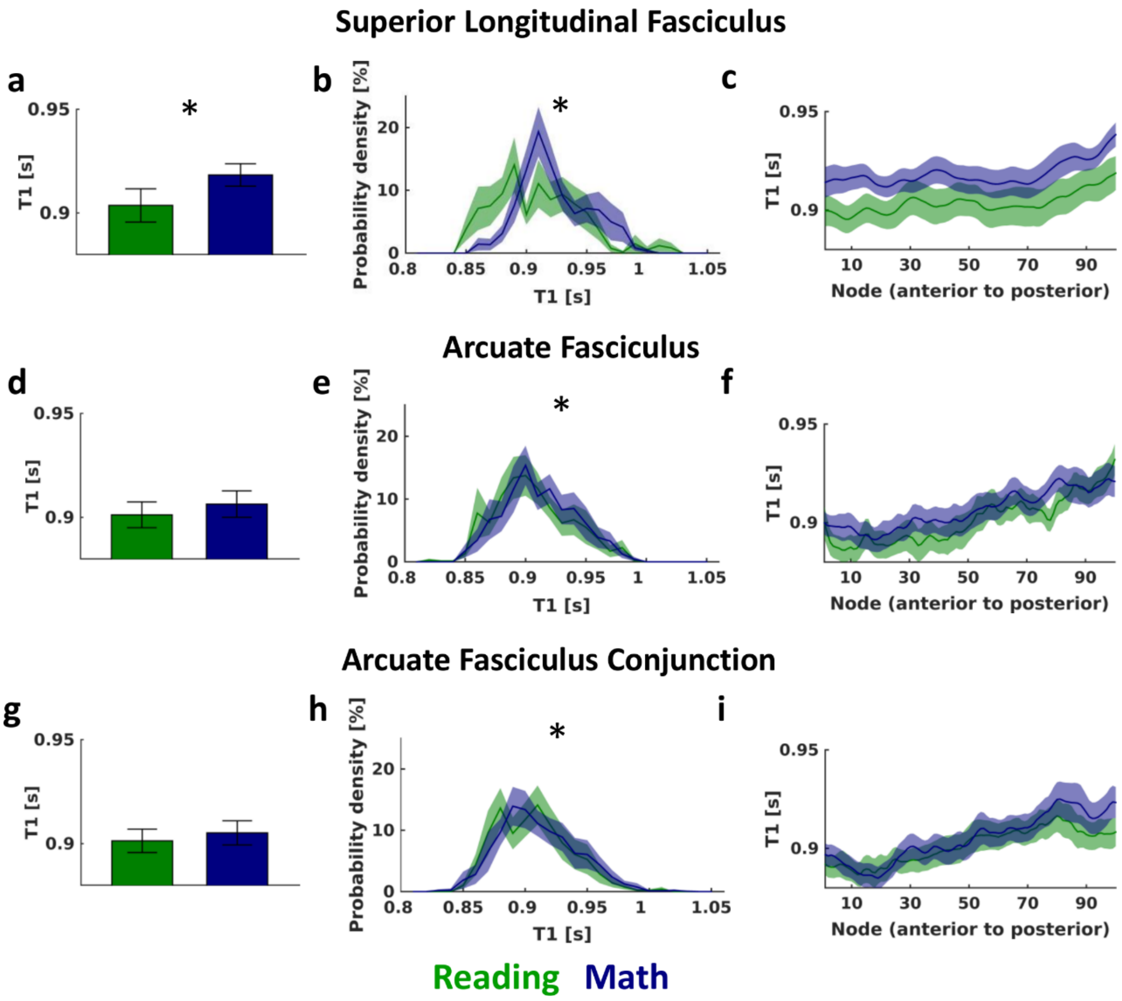


**Supplementary Fig. 23. Same as Fig. 5, but for tracts identified using fROI to fROI tractography. (a-c)**: $T_1$ measurements for SLF tracts connecting IFG and SMGr in the reading network (green) and PCS and SMGm in the math network (blue). **(d-f)**: $T_1$ measurements for AF tracts connecting the IFG and STS in the reading network (green) and the PCS and ITG in the math network (blue). **(g-i)**: $T_1$ measurements for AF tracts connecting the lOTC conjunction fROI with the IFG in the reading network (green) and the PCS in the math network (blue). Left **(a,d,g)**: Average $T_1$ for reading- and math-related tracts in the SLF and the AF. Bar graph shows mean across subjects ± SEM. *: $T_1$ for math and reading related tracts differs significantly, $p<0.05$. Middle **(b,e,h)**: Distribution of $T_1$ values across all tracts. Distributions were calculated within each subject and node; the plot shows the mean across all nodes ± SEM. *: Distributions differ significantly, $p<0.05$. Right **(c,f,i)**: Average $T_1$ for reading- and math-related tracts along the SLF and the AF. Line graph shows mean across subjects ± SEM. *Abbreviations:* IFG=inferior frontal gyrus, PCS=precentral sulcus, SMGr=reading fROI in supramarginal gyrus, SMGm=math fROI in supramarginal gyrus, STS=superior temporal sulcus, ITG=inferior temporal gyrus, lOTC=lateral occipito-temporal cortex, SLF=superior longitudinal fasciculus, AF=arcuate fasciculus.
